# Gamma (γ)-radiation stress response of the cyanobacterium *Anabaena* sp. PCC7120: Regulatory role of LexA and photophysiological changes

**DOI:** 10.1101/2022.07.04.498681

**Authors:** Akanksha Srivastava, Arvind Kumar, Subhankar Biswas, Rajender Kumar, Vaibhav Srivastava, Hema Rajaram, Yogesh Mishra

## Abstract

High radioresistance of the cyanobacterium, *Anabaena* sp. PCC7120 has been attributed to efficient DNA repair, protein recycling, and oxidative stress management. However, the regulatory network involved in these batteries of responses remains unexplored. In the present study, the role of a global regulator, LexA in modulating gamma (γ)-radiation stress response of *Anabaena* was investigated. Comparison of the cytosolic proteome profiles upon γ-radiation in recombinant *Anabaena* strains, AnpAM (vector-control) and An*lexA*^+^ (LexA-overexpressing), revealed 41 differentially accumulated proteins, corresponding to 29 distinct proteins. LexA was found to be involved in the regulation of 27 of the corresponding genes based on the presence of AnLexA-Box, EMSA, and/or qRT-PCR studies. The majority of the regulated genes were found to be involved in C-assimilation either through photosynthesis or C-catabolism and oxidative stress alleviation. Photosynthesis, measured in terms of PSII photophysiological parameters and thylakoid membrane proteome was found to be affected by γ-radiation in both AnpAM and An*lexA*^+^ cells, with LexA affecting them even under control growth conditions. Thus, LexA functioned as one of the transcriptional regulators involved in modulating γ-radiation stress response in *Anabaena*. This study could pave the way for a deeper understanding of the regulation of γ-radiation-responsive genes in cyanobacteria at large.

**Highlights:**

- γ-radiation alters PSII photophysiology and thylakoid proteome profile in *Anabaena*.
- LexA modulates the cytosolic and thylakoid proteome of *Anabaena* under γ-radiation.
- LexA functions as one of the regulators of radiation-responsive genes in *Anabaena*.

## 1. Introduction

Radiation has co-existed since the beginning of earth, though the intensity and sources have varied over the years. However, the evolution of radiation-resistant microbes has continued from ancient cyanobacteria (Billi et al., 2000; Singh et al., 2013; Badri et al., 2015; Verseux et al., 2017) and Archaeon *Thermococcus gammatolerans* (Zivanovic et al., 2009) to more recent *Deinococcus radiodurans* (Cox and Battista, 2005) and *Halobacterium* sp. (Kottemann et al., 2005) that can withstand radiation doses over 5.0 kGy. It has long been considered that the DNA is the primary target for γ-radiation damage, and that the high radioresistance in few organisms is due to their unique DNA protection and repair mechanisms. Growth phase-dependent analysis of DNA repair revealed faster stitching of shattered chromosome in the stationary phase in *T. gammatolerans*, resulting in growth-phase independent cell survival against radiation in this organism contrary to that observed in other bacteria (Tapias et al., 2009). In addition to DNA damage, γ-radiation has also been shown to induce significant morpho-ultrastructural, genetic, biochemical, physiological, and proteomic changes across organisms (Strydom et al., 1991; Maity et al., 2005; Agarwal et al., 2008; Caillet et al., 2009; Jan et al., 2012; Badri et al., 2015; Van Hoeck et al., 2015; Singh and Apte, 2018; Pradhan et al., 2020). Exposure to γ-radiation up to 2.5 kGy stimulated the content of phenols, proline, soluble protein, minerals (N, P, Na, K, Ca, Mg, and Fe), and vitamins (A, K, and B group), as well as the activities of N-assimilating and antioxidant enzymes in the cyanobacterium, *Arthrospira platensis* (Shabana et al., 2017). And higher doses up to 5 kGy of γ-radiation reduced the expression of genes involved in photosynthesis, carbon assimilation, and pigment synthesis, while inducing the expression of genes involved in cellular protection, detoxification, and repair in *Arthrospira* sp. PCC8005 (Badri et al., 2015). A highly radioresistant cyanobacterium, *Chroococcidiopsis* spp. was found to survive high doses (11.59 kGy) of γ-radiation with no noticeable damage to DNA or plasma membranes (Verseux et al., 2017).

Proteomic changes induced during exposure to γ-radiation and during post-irradiation recovery have been reported across organisms. In Archaeon, *T. gammatolerans*, a specific feature of proteome changes was the increase in cellular detoxification proteins (Zivanovic et al., 2009). In *D*. *radiodurans*, proteomic changes was considered to be the primary means of conferring radiotolerance rather than only DNA repair (Daly et al., 2007; Krisko and Radman, 2010), with protein recycling observed for proteins from varied functional groups (Zhang et al., 2005; Basu and Apte, 2012). This implied that induction of these stress-responsive proteins assists in the repair of damaged cellular components, including the DNA, and eventually restoration of other impaired metabolic functions. Among cyanobacteria, *Anabaena* sp. PCC7120 (hereinafter referred to as *Anabaena*) responded to the LD_50_ dose of γ-radiation (6 kGy) through a flux of proteomic changes that included decreased abundance of proteins related with photosynthesis, respiration, and carbon and nitrogen assimilation, and increased abundance of proteins related with reactive oxygen species (ROS) detoxifiers, chaperones, and proteases (Singh et al., 2013; Singh and Apte, 2018). Irradiated *Anabaena* cells exhibit 2-3 days of lag period before they show growth recovery, which is matched by the restoration of intact DNA, as well as the proteome profile (Singh et al., 2013). In addition to the expected changes in proteome profile due to changes in gene expression during post-irradiation recovery (PIR), several proteins exhibited increased abundance immediately after irradiation (Singh et al., 2013; Singh and Apte, 2018). This is suggestive of regulation of gene expression during exposure to γ-radiation, in addition to that during PIR. However, the regulatory mechanism(s) involved in these proteomic changes have not been identified in cyanobacteria.

LexA is a well-known transcriptional repressor of SOS genes in *Escherichia coli* and several other heterotrophic bacteria (Eril et al., 2007). However, in cyanobacteria, it exhibits a varied and global cellular response. In a unicellular cyanobacterium, *Synechocystis* sp. PCC6803 (hereinafter referred to as *Synechocystis*), it has been shown to be crucial for survival under carbon starvation (Domain et al., 2004). It was found to regulate a variety of genes, such as those coding for bidirectional hydrogenase (Gutekunst et al., 2005; Oliveira and Lindblad, 2005; Antal et al., 2006), RNA helicase (Patterson-Fortin and Owttrim, 2008), bicarbonate transporter gene (Lieman-Hurwitz et al., 2009), fatty acid biosynthesis genes (Kizawa et al., 2017), and salt stress-inducible genes (Takashima et al., 2020) in *Synechocystis* and *hyp* operon in *Lyngbya majiscula* (Leitao et al., 2006; Ferreira et al., 2007). In the filamentous cyanobacteria, it was also found to regulate bidirectional hydrogenase (Sjoholm et al., 2007), *lexA*, *recA*, and *ssb* (Li et al., 2010; Kirti et al., 2017; Kumar et al., 2018), *katB* and *sodA* (Kumar et al., 2018), *sbcC* and *sbcD* (Pandey et al., 2018), several photosynthetic genes (Srivastava et al., 2022), in *Anabaena* and the *hoxW* gene in *Nostoc punctiforme* ATCC29133 (Devine et al., 2009). While the regulation by LexA in cyanobacteria has been shown to be through its binding to the LexA-Box-like element, the sequence corresponding to this box has varied (Mazón et al., 2004; Sjoholm et al., 2007; Kumar et al., 2018).

Several genes coding for DNA repair/metabolism proteins (Rajaram et al., 2020) and proteins involved in oxidative stress alleviation (Kumar et al., 2018), have been shown to possess the AnLexA-Box in *Anabaena*. However, low expression levels of the DNA repair proteins in *Anabaena* resulted in them not being picked up during proteomic studies of irradiated cultures and during post-irradiation recovery (Singh and Apte, 2018). Due to the observed global regulatory role of LexA in stress response and decreased radioresistance of *Anabaena* cells overexpressing LexA (An*lexA*^+^) (Kumar et al., 2018), it was envisaged that the LexA may be involved in modulating protein expression during γ-radiation in *Anabaena*. The level of the LexA protein in An*lexA*^+^ cells was ∼10-fold higher than in AnpAM cells, and the cells exhibited no change in morphology or growth under normal growth conditions upon LexA overexpression (Kumar, 2019 Ph.D. Thesis). However, upon exposure to abiotic stresses, it exhibited increased growth in the presence of DCMU upon C-starvation, but decreased tolerance to γ-radiation, heavy metal, and oxidative stress (Kumar et al., 2018). In this study, we focussed on obtaining insight into regulation of the changes in protein expression occurring in *Anabaena* during γ-radiation, i.e., before the cells undergo post-irradiation recovery, as the cellular integrity at that stage would determine its ability to proceed for recovery. For this purpose, the LD_50_ dose of γ-radiation (6 kGy) was taken, as earlier studies had shown significant proteomic changes on this dose (Singh et al., 2013). Efforts at obtaining *lexA* knock out mutant in *Anabaena* were unsuccessful as the reversion to wild type was very high (Kumar et al., 2018). Our observation during the tracking of the neomycin cassette was that when the *lexA* gene was replaced by the neo^R^ cassette in more than 20% of the cells of the filament, the cells died. Even those with 20% were highly photobleached indicating extreme loss of photosynthesis. This was not surprising because LexA has been shown to up-regulate RbcL and down-regulate C-catabolism genes (Kumar et al., 2018), as a result, in the absence of LexA, the C-reserve would be decreasing, and with the added involvement of LexA in regulating photosynthetic electron transport chain (pETC) genes (Srivastava et al., 2022), it probably rendered the *lexA* deletion mutant cells non-viable. In the absence of a mutant strain due to its non-viability, use of a strain overexpressing the protein can provide insights into the physiological role of the gene in that organism instead of being dependent on a heterologous system. Thus the LexA-overexpressing recombinant *Anabaena* strain, An*lexA*^+^ was used for the studies. Since, the An*lexA*^+^ cells (Kumar et al., 2018) are grown in the presence of the antibiotic neomycin, vector control AnpAM harbouring the pAM1956 plasmid (Rajaram and Apte, 2010) was used as a control in place of the wild type *Anabaena*. Regulation of genes coding for some of the identified proteins either by γ-radiation or by LexA or both was confirmed by quantitative reverse transcriptase-PCR (qRT-PCR), presence of AnLexA-boxes, and electrophoretic mobility shift assay (EMSA) studies to show binding of LexA to the upstream regions of genes predicted to be regulated by LexA. Photophysiological changes were also analysed to assess the damage to the photosynthetic complexes upon γ-irradiation in conjunction with LexA.

## 2. Materials and methods

### 2.1 Strains used, growth and stress conditions

Axenic cultures of AnpAM (Rajaram and Apte, 2010) and An*lexA*^+^ (Kumar et al., 2018) were grown in combined nitrogen-added BG-11 medium, pH 7.2 (Castenholz, 1988) with 10 μg neomycin mL^−1^ at 25 ± 2 °C in continuous day light fluorescent tubes (40W, Philips, Bangalore, India) emitting 72 μmol photon m^−2^ s^−1^ photosynthetic photon flux density (PPFD). Three-day-old (exponential phase) AnpAM and An*lexA*^+^ cultures, concentrated to 10 μg Chl *a* mL^−1^ were either exposed to 6 kGy ^60^Co γ-radiation, dose rate 4.5 kGy h^−1^ (gamma cell 5000 irradiation unit, BARC) (IR), or kept in the dark under similar conditions but without exposure to irradiation (sham-irradiated control, SIC). Cells were irradiated in glass vials kept in water to ensure minimum increase in heat due to radiation. The duration of γ-irradiation was about 1 h 20 min during which the cells remained in the dark. For the entire period that the treated cells remained in the dark, the corresponding control cultures were likewise kept in the dark (sham irradiated control, SIC) to ensure that only changes due to irradiation are accounted for and not those occurring due to lack of illumination in the irradiated cells. The control and irradiated cells were harvested immediately after γ-irradiation or SIC without exposing them to light, to assess the change in transcript and protein profile during γ-radiation. An*lexA*^+^ cells were monitored microscopically for expression of GFP and for LexA in protein gels to ensure that only fully segregated recombinant An*lexA*^+^ cells are used for the experiments. Cell viability was checked immediately after irradiation for both AnpAM and An*lexA*^+^ cells (data not shown) and were found to conform to earlier results (Kumar et al., 2018).

### 2.2 Chlorophyll a fluorescence measurement

Rapid light curves (RLCs) of recombinant *Anabaena* strains were monitored at room temperature using Dual-PAM-100 (Heinz Walz GmbH, Effeltrich, Germany). There were 11 photosynthetically active radiation (PAR) gradients (0, 30, 37, 46, 77, 119, 150, 240, 363, 555, and 849 µmol m^−2^ s^−1^) used, each with a 20 s irradiation time. The saturation pulse used in the RLCs measurement was 3000 μmol photons m^−2^ s^−1^ for 0.6 s. All relevant PSII photophysiological parameters, including the effective quantum yield of PSII [Y(II)], the puddle model-based coefficient of photochemical quenching (qP), the lake model-based coefficient of photochemical quenching (qL), the electron transport rate through PSII [ETR(II)], and the quantum yield of non-regulated heat dissipation in PSII [Y(NO)], were automatically calculated by the Dual-PAM software.

### 2.3 Blue native (BN)-PAGE

For thylakoid membrane (TM) proteome analysis of recombinant *Anabaena* strains, TMs were isolated and subjected to BN-PAGE, as described previously (Srivastava et al., 2021a,b). The experiment was repeated three times and a representative image included. The quantification of band signal in BN-PAGE gel was done using ImageJ software (http://rsb.info.nih.gov/ij) and background correction applied. To assess change in levels of different complexes due to LexA overexpression, band intensities of An*lexA*^+^, SIC were compared with those of AnpAM, SIC. In contrast, for assessing the effect of γ-radiation, band intensities of AnpAM, IR was compared with that of AnpAM, SIC and of An*lexA*^+^, IR with that of An*lexA*^+^, SIC.

### 2.4 Isoelectric focusing (IEF)-SDS PAGE

For cytosolic proteome analysis of recombinant *Anabaena* strains, total cell proteins were extracted, precipitated, and solubilized, as described previously (Kaur et al., 2019). Protein separation through IEF followed by SDS-PAGE was performed, as described earlier (Srivastava et al., 2021b). The intensity of each spot in each of the four samples was measured and the spot intensities from AnpAM, IR and An*lexA*^+^, SIC were normalized by dividing with the corresponding spot intensity of AnpAM, SIC, and the spot intensities from An*lexA*^+^, IR were normalized by the corresponding spot intensity of An*lexA*^+^, SIC. The protein spots with statistically significant and reproducible changes (>1.5 fold, *p* < 0.05) were designated as <u>D</u>ifferentially <u>A</u>ccumulated <u>P</u>rotein (DAP) spots. These DAP spots were manually cut from the gel using a sterile scalpel and processed for mass spectrometry, as previously described (Srivastava et al., 2021a,b). Peptides with mascot scores greater than the statistical significance criterion (*p* < 0.05) were chosen.

### 2.5 In silico prediction of AnLexA-box in genes encoding identified cytosolic DAPs

The upstream regions of 29 genes coding for identified differentially accumulated cytosolic proteins were bioinformatically analysed for the presence of a bacterial promoter using the BPROM software (http://www.softberry.com/berry.phtml?topic) (Solovyev and Salamov, 2011), and the presence of AnLexA-Box, as defined earlier (Kumar et al., 2018; Srivastava et al., 2022).

### 2.6 Transcript analysis

Total RNA was extracted from the recombinant *Anabaena* strains and used for cDNA synthesis, as described previously (Srivastava et al., 2021a). Primer designing for the 11 selected genes was carried out using Primer-BLAST (NCBI), listed in Supplementary Table S1. Transcript expression analysis was done through qRT-PCR, as described earlier (Srivastava et al., 2021a), using CFX-96 Real-time PCR Detection System (Bio-Rad, USA). Normalization of the transcript levels was carried out with respect to *rnpA* and *secA*, and the fold change was calculated using the 2^−ΔΔCt^ method. Significant difference in transcript levels was considered only when the fold change was ≥ 1.3 and *p* < 0.05 for high confidence.

**Table 1.**
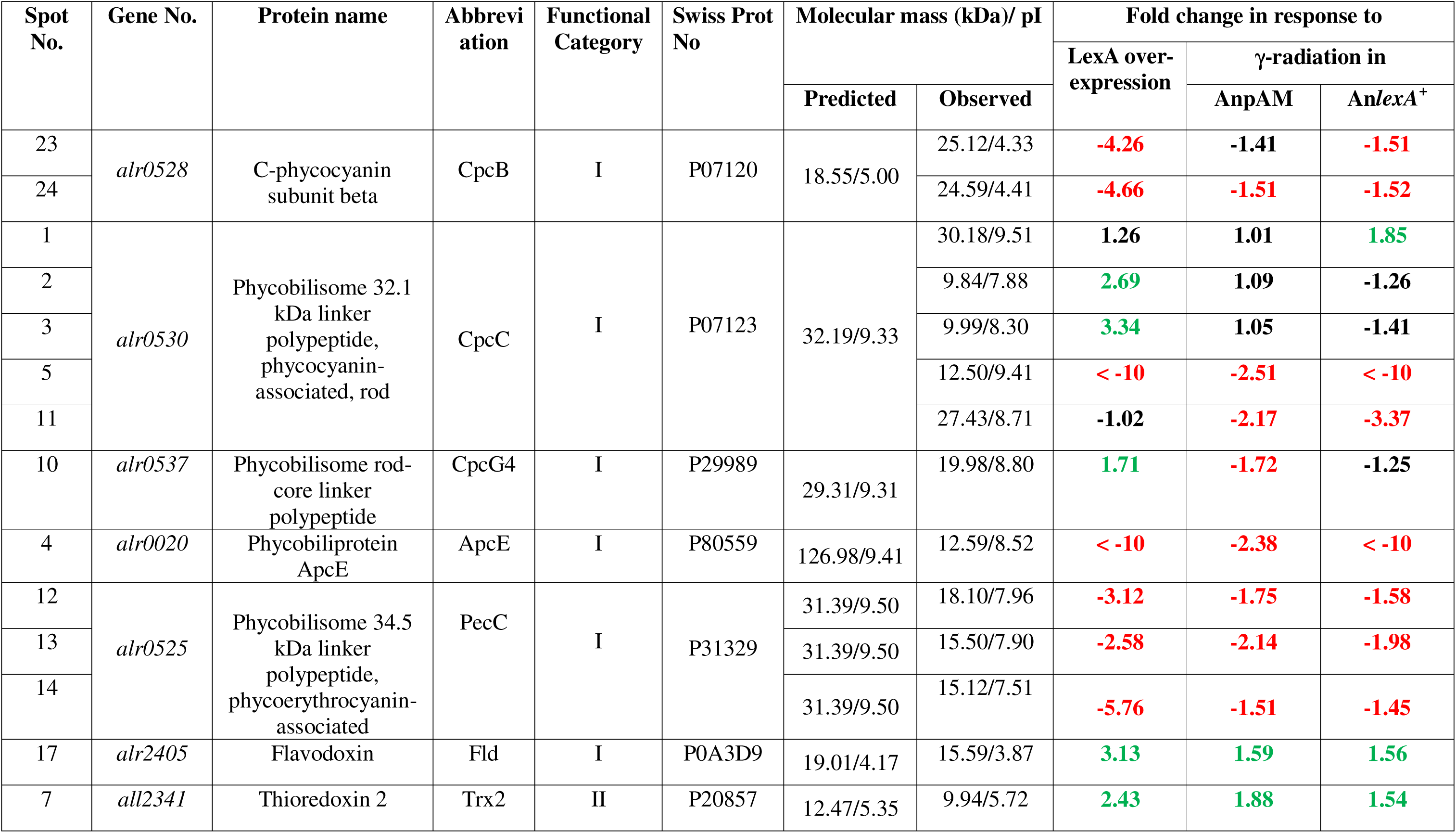
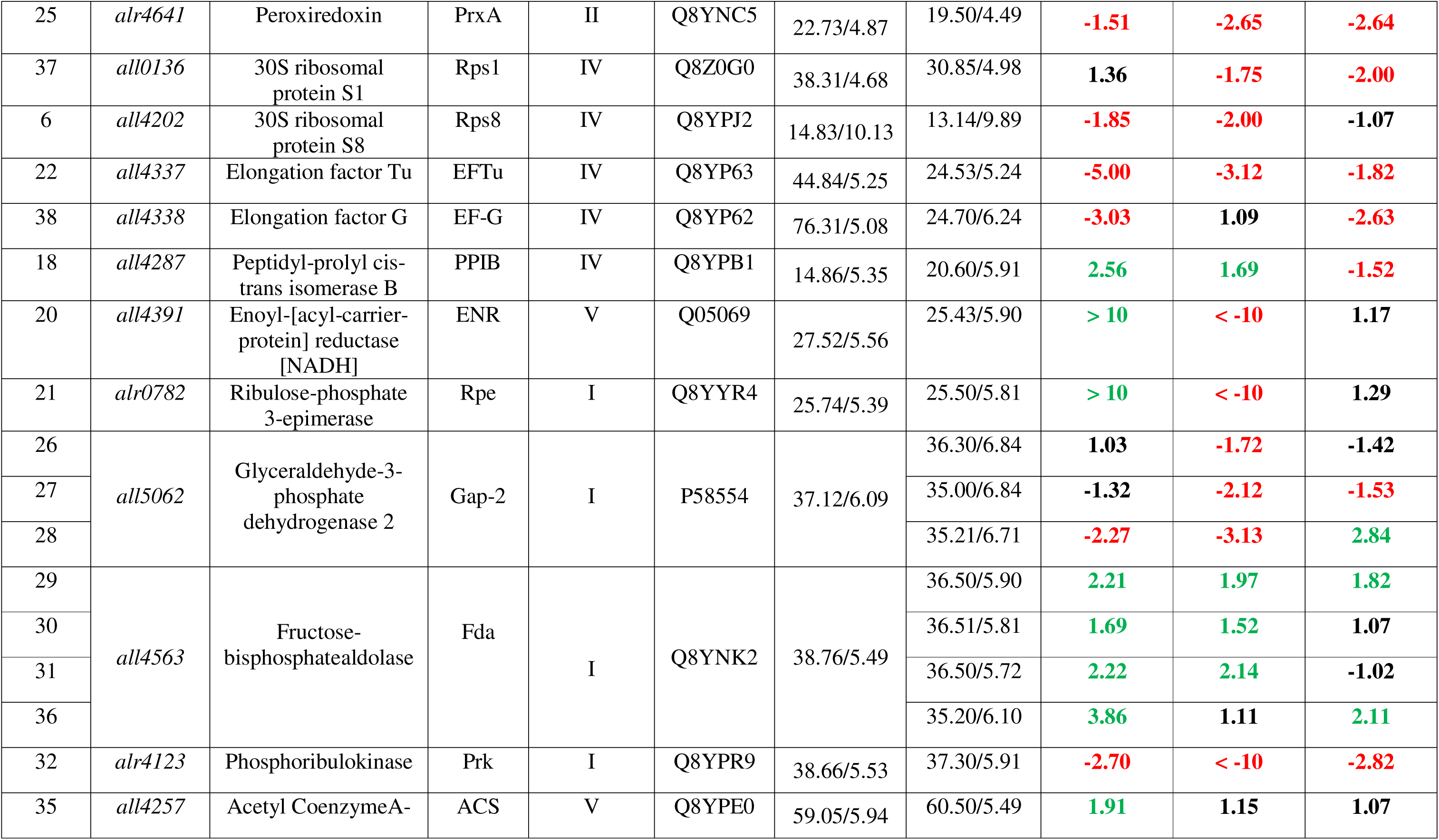
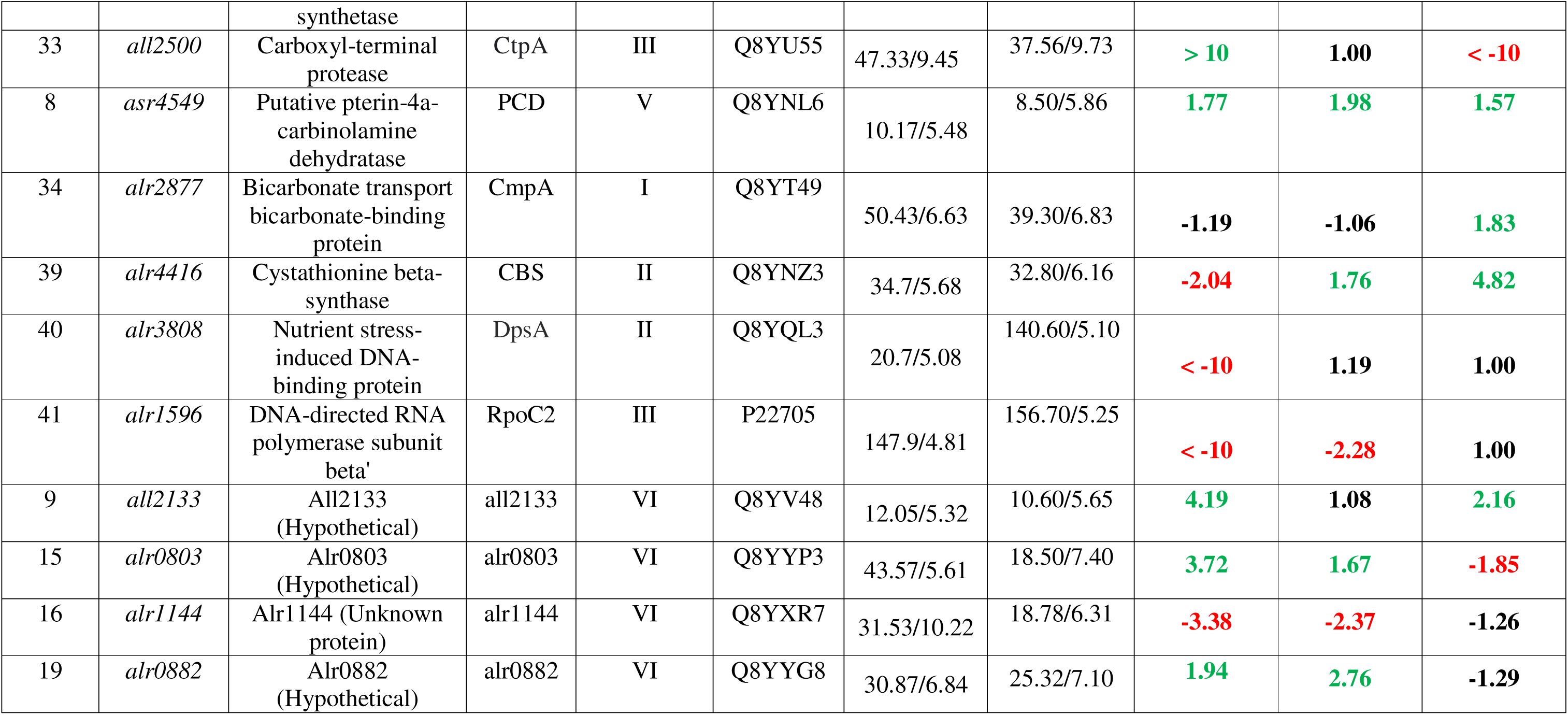
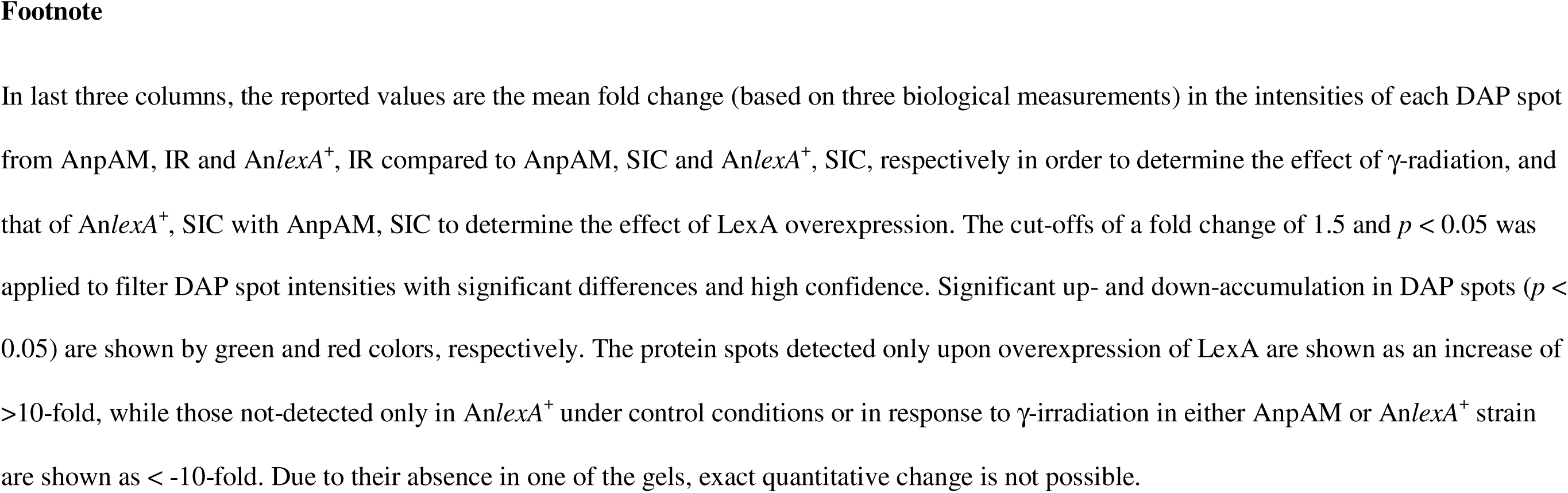
Identification details of 41 DAP spots detected in cytosolic proteome profiling of recombinant *Anabaena* strains AnpAM (vector control) and An*lexA*^+^ (LexA-overexpressing), under sham-irradiated control (SIC) and γ-irradiated (IR) conditions, as well as mean fold changes in their intensities

### 2.7 Electrophoretic mobility shift assay (EMSA)

A 200–300 bp upstream region of five selected genes for EMSA were amplified using the primers provided in Supplementary Table S2. Binding of LexA to DNA fragments was assessed by EMSA using the specified concentrations of purified *Anabaena* LexA and DNA fragments, as described earlier (Kirti et al., 2017), followed by staining with SYBR Green I, visualisation using a UV transilluminator, and quantification of band intensities using ImageJ software (http://rsb.info.nih.gov/ij).

### 2.8 Statistical analysis

Data were presented as the mean ± standard error (SE) of at least three biological replicates. To assess the significant differences in recombinant *Anabaena* strains, data were statistically examined using Student’s t-test with Tukey’s post-hoc test at a significant level of *p* < 0.05. The differences in DAPs from IEF-SDS PAGE were examined by principal component analysis (PCA) using BioDiversity Pro 2.0 (McAleece et al., 1997).

## 3. Results

In order to assess the role of LexA in modulating γ-radiation response, the proteomic changes occurring in irradiated An*lexA*^+^ cells with respect to its sham-irradiated control (SIC) was compared to the proteomic changes in irradiated AnpAM cells with respect to its SIC. This was further validated through qRT-PCR and EMSA analyses. In this process, we also identified a few proteins showing change in abundance upon overexpression of LexA under control conditions (An*lexA*^+^, SIC *vs* AnpAM, SIC), and upon irradiation when LexA is not overexpressed (AnpAM, IR *vs* AnpAM, SIC), which were not reported earlier (Kumar et al., 2018; Singh and Apte, 2018). The physiological implications were analysed through study of PSII photophysiological parameters and thylakoid membrane complexes.

### 3.1 Effect of LexA overexpression on cytosolic proteome changes in Anabaena in response to **γ**-radiation stress

Fig. 1 illustrates four representative IEF-SDS PAGE gels for the four sets of samples (AnpAM, SIC; AnpAM, IR; An*lexA*^+^, SIC; and An*lexA*^+^, IR). Within the molecular mass range of 8–240 kDa and isoelectric point range of 3–10, 254, 249, 261, and 269 reproducible protein spots were detected respectively in AnpAM, SIC; AnpAM, IR; An*lexA*^+^, SIC; and An*lexA*^+^, IR, of which 168 could be matched in all four gels and their triplicates. Among these, the 41 differentially accumulated protein (DAP) spots (>1.5 fold, *p* < 0. 05) in response to either γ-radiation or LexA overexpression or both, were identified and listed in Table 1.

**Fig. 1.**
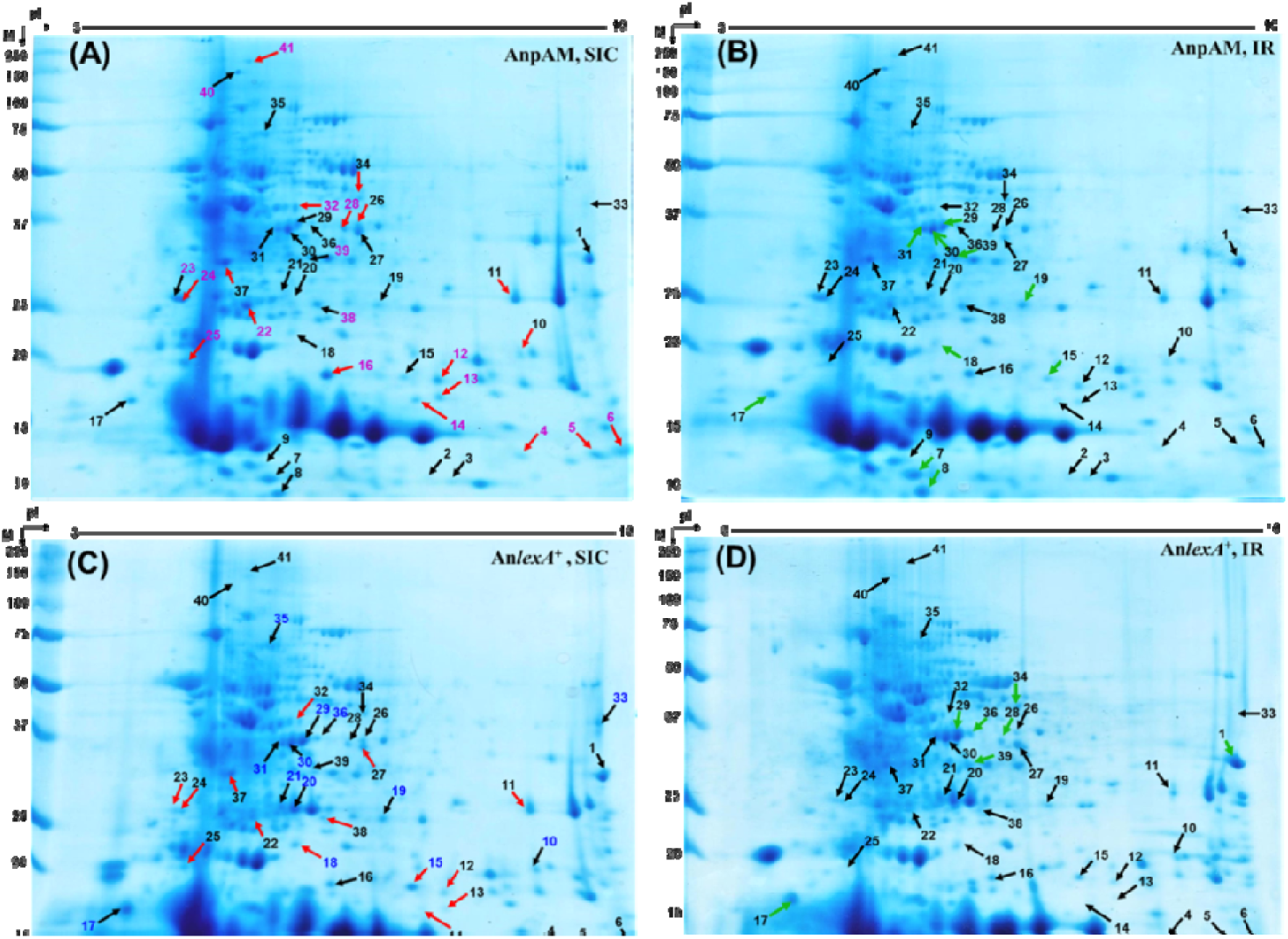
Cytosolic proteome profiling corresponding to (A and C) sham-irradiated control (SIC) and (B and D) 6 kGy ^60^Co γ-irradiated (IR) recombinant *Anabaena* strains (A and B) AnpAM (vector control) and (C and D) An*lexA*^+^ (LexA-overexpressing) Identified proteins are labelled with their respective spot numbers in the representative images. The red arrows in (A) and (C) correspond to proteins showing lower abundance, and the green arrows in (B) and (D) correspond to proteins showing higher abundance upon IR. The magenta (or blue) spot numbers correspond to proteins showing lower (or higher) abundance upon LexA overexpression. Molecular masses of protein size markers are indicated in the left in kDa and pI is shown on the top.

Of the 41 DAP spots, γ-radiation resulted in differential abundance of 28 protein spots in AnpAM and 25 in An*lexA*^+^, 15 of which were common (Supplementary Table S3). However, the fold change of these DAP spots differed in AnpAM and An*lexA*^+^ (Table 1). In response to LexA overexpression under control conditions (SIC), 18 protein spots showed increased abundance, of which 8 showed increased abundance in γ-irradiated AnpAM and 6 in γ-irradiated An*lexA*^+^, three of them being common to both (Supplementary Table S3). Decreased abundance was observed for 17 protein spots in An*lexA*^+^, SIC compared to AnpAM, SIC, of which 12 showed decreased abundance in AnpAM, IR and 9 in An*lexA*^+^ IR (Supplementary Table S3), which is suggestive of involvement of LexA in regulating some of the γ-radiation responsive genes.

PCA demonstrated a distinct separation among AnpAM, SIC; AnpAM, IR; An*lexA*^+^, SIC; and An*lexA*^+^ (Fig. 2A), IR indicating that the cytosolic proteomes of these four samples were significantly different. Furthermore, in PCA plots (Fig. 2A), An*lexA*^+^, SIC and IR were clustered differently than AnpAM, SIC and IR, indicating that LexA overexpression does affect the cytosolic proteome of *Anabaena* under both the conditions. The PCA of 41 DAP spots revealed that few protein spots were segregated from others and these were deemed probable outliers and prioritized for mass-spectrometry investigation (Fig. 2B).

**Fig. 2.**
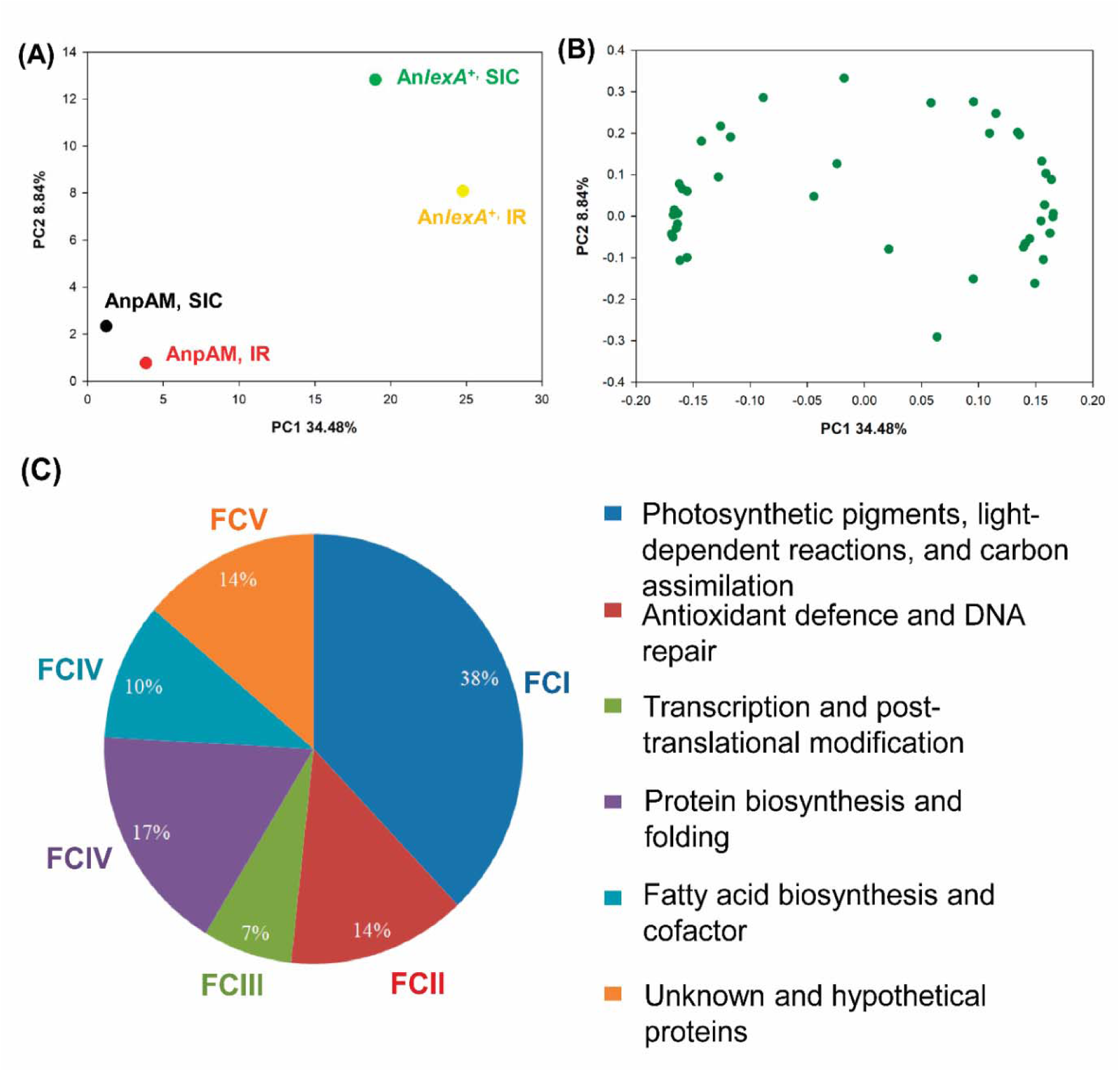
(A) Principal component analysis (PCA) of sham-irradiated control (SIC) and γ-irradiated (IR) recombinant *Anabaena* strains AnpAM (vector control) and An*lexA*^+^ (LexA-overexpressing) using BioDiversity Pro 2.0 (B) PCA of and (C) functional classification of 41 DAP spots identified from aforementioned samples, namely AnpAM, SIC; AnpAM, IR; An*lexA*^+^, SIC; and An*lexA*^+^, IR.

The 41 DAP spots corresponded to 29 proteins, as some of the proteins were detected as multiple spots on the gel (Table 1). Of the 41 DAP spots, increased abundance was observed for spot nos. 2, 3, 20 and 21 exclusively upon overexpression of LexA under control conditions, while that of spot no. 40 decreased (Table 1, Supplementary Table S3). Of these spot nos. 2 and 3, corresponded to CpcC protein, which was detected as multiple spots (Table 1), while spot no. 20, 21 and 40 corresponded to ENR, RpE, and DpsA (Table 1). All other protein spots showed change in abundance in response to γ-radiation in either AnpAM or An*lexA*^+^ (Table 1, Supplementary Table S3).

To obtain insights into the protein families most affected by IR in the absence or presence of constitutively expressed high levels of LexA, the identified DAPs were classified into six functional categories (FCs) (Fig. 2C) namely, (i) FCI- photosynthetic pigments, light-dependent reactions, and carbon assimilation, (ii) FCII- antioxidant defence and DNA repair, (iii) FCIII- transcription and post-translational modification, (iv) FCIV- protein biosynthesis and folding, (v) FCV- fatty acid biosynthesis and cofactor, and (vi) FCVI- unknown and hypothetical proteins. Of these, maximal changes were observed in FC1 category. The functional category of each identified protein spot is indicated in Table 1.

### 3.2 Assessment of transcriptional regulation of genes encoding the identified cytosolic DAPs

Molecular mass of some of the protein spots did not match with the actual molecular mass of the corresponding identified protein (Fig. 1, Table 1), which has been observed earlier as well for *Anabaena* proteome (Singh et al., 2013; Kumar et al., 2018; Singh and Apte, 2018). The proteome profile of An*lexA*^+^, SIC (Fig. 1C) differed for some of the spots in terms of fold change as compared to that reported for An*lexA*^+^ earlier (Kumar et al., 2018), because the SIC were kept in dark for the duration of the irradiation experiment and the pre- and post-processing time, which was around 2 h. Hence, to confirm if the genes encoding proteins corresponding to these spots were actually regulated by LexA, they were analysed bioinformatically for the presence of AnLexA-Box in their upstream regulatory region, followed by EMSA studies to assess binding of LexA to a few of the predicted AnLexA-boxes. AnLexA-box was detected in the upstream regulatory region of 27 genes including two, namely *rps8* and *all2133*, wherein it was present upstream of their gene preceding (Table 2). The two genes, which have no AnLexA-Box, are *fus* (EF-G) and *tufA* (EF-Tu) and they function as an operon (Table 2). The distance between the predicted AnLexA-Box upstream of *prk* and *alr0803* genes at the predicted -10 region was found to be over 100 bases (Table 2). Of these, regulation of 4 genes, corresponding to 9 protein spots, namely *cpcB*, *cpcC*, *apcE*, and *prxA* by LexA was shown earlier (Kumar et al., 2018). Further, the binding of purified *Anabaena* LexA was tested for upstream region of 5 genes, namely *pecC*, *all4287* (PPIB), *gap-2*, *prk*, and *alr0803* (Fig. 3).

**Fig. 3.**
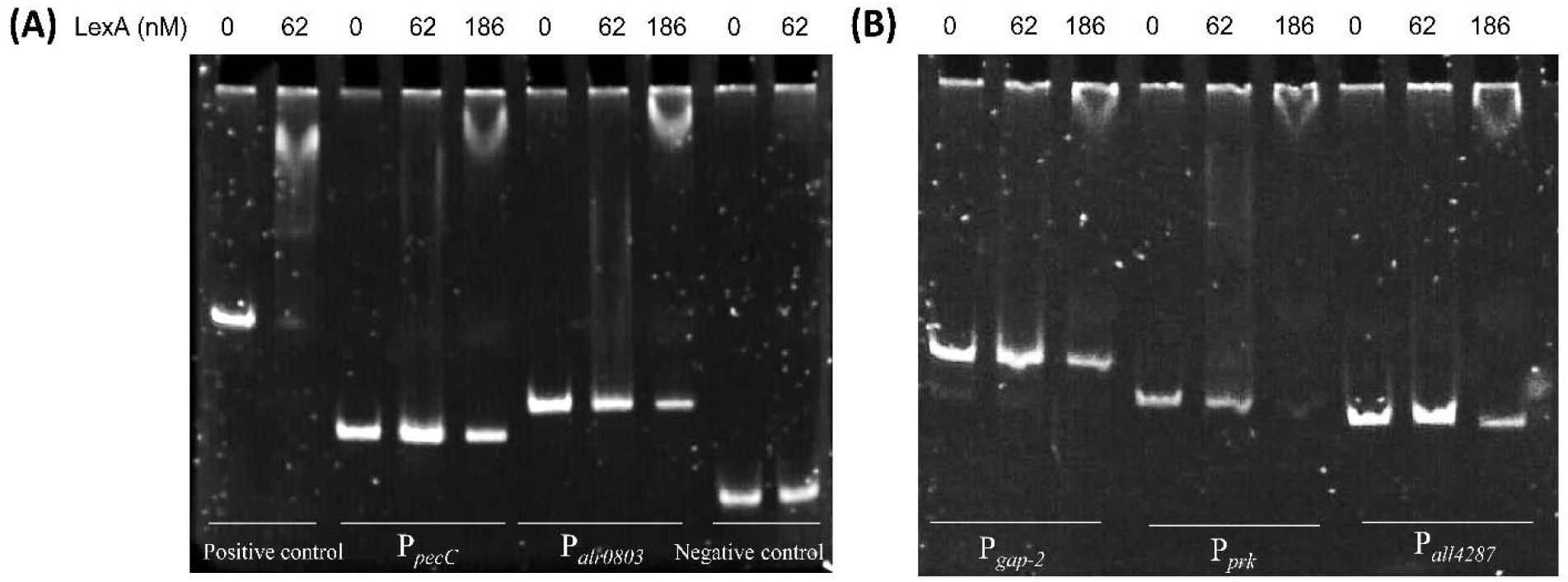
*In vitro* interaction of *Anabaena* LexA protein with DNA fragments using electrophoretic mobility shift assay (EMSA). DNA fragments (30 ng) corresponding to upstream regions of (A) *lexA* (positive control), *pecC*, *alr0803*, and negative control (DNA fragment with no AnLexA-Box) and (B) *gap-2*, *prk*, and *all4287* were incubated with 0, 62 and/or 186 nM LexA and electrophoretically separated on native 8% polyacrylamide gel followed by staining with SYBR Green I.

**Table 2.**
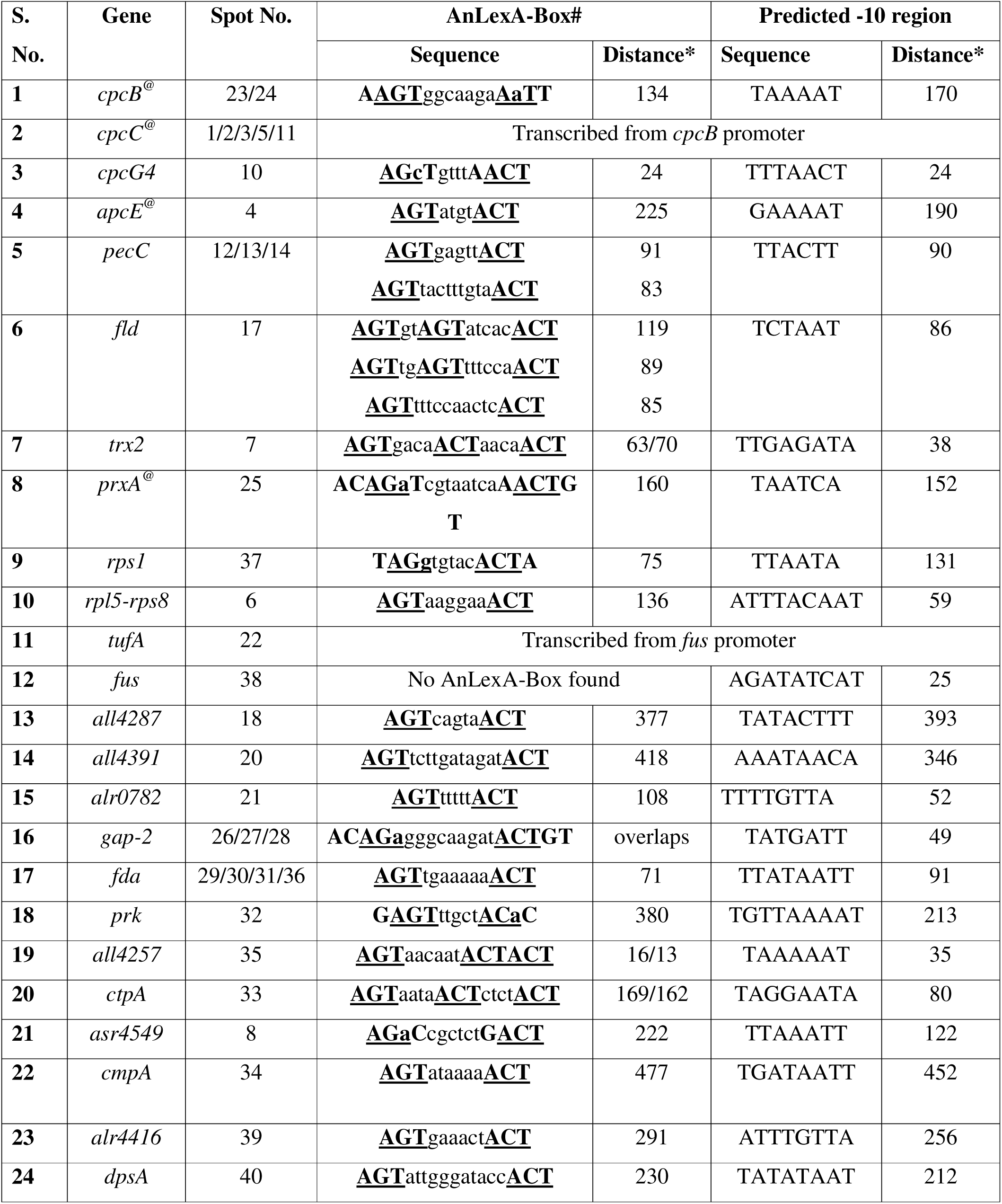
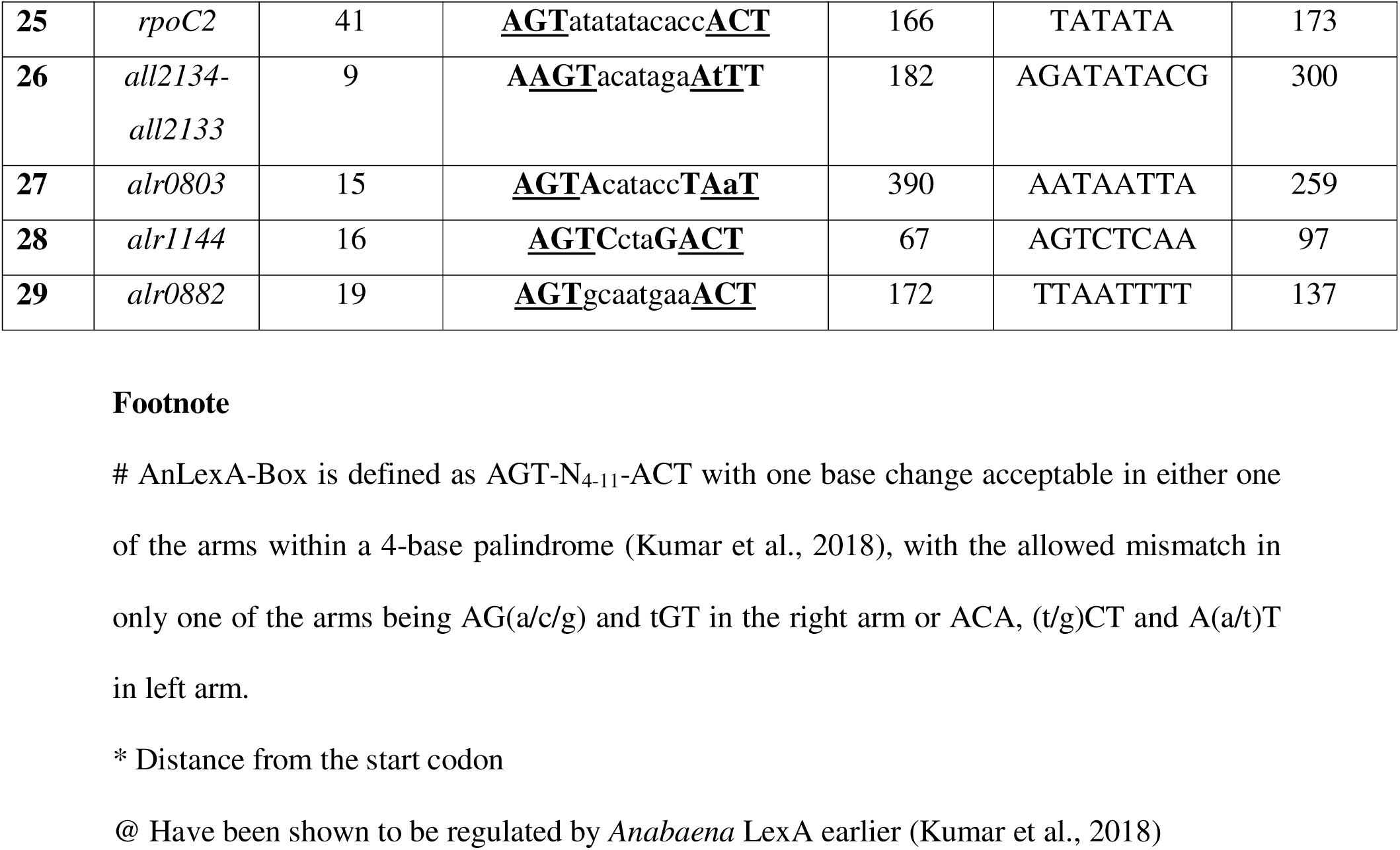
*In silico* AnLexA-Box prediction in the identified DAPs encoding genes

Since, earlier results had shown the binding affinity of LexA to be 18-96 nM, only two specific concentrations of LexA (62 nM and 186 nM) were used for binding studies. Shift in mobility was observed for all the five promoter regions tested with ∼20-30% binding with 62 nM LexA and over 70% binding in the presence of 186 nM LexA, while no binding was observed for negative control DNA fragment which lacked the AnLexA-Box (Fig. 3).

For analysing transcriptional regulation by IR or LexA or both, qRT-PCR was carried out for 11 selected genes (corresponding to 18 protein spots), encoding the proteins PecC, Rps1, EF-Tu, EF-G, PPIB, Gap-2, Fda, Prk, CtpA, CmpA, and Alr0803 (Fig. 4). Comparison of proteomic and transcript data upon LexA overexpression [An*lexA*^+^, SIC compared with AnpAM, SIC] revealed (i) complete agreement in the data corresponding to *pecC* and *gap-2*, both the protein spots and transcript being down-regulated, (ii) increase in abundance of protein spots corresponding to Fda, PPIB, and CtpA but decreased transcript levels, (iii) increased abundance of Alr0803 but no change in transcript levels, (iv) decreased abundance of Prk, EF-Tu, and EF-G but no change in transcript levels, and (v) no change in abundance of CmpA and Rps1, but decreased transcript levels (Fig. 4). This clearly indicates that of these 11 genes, only 7 namely *pecC*, *gap-2*, *fda*, *all4287*, *ctpA*, *cmpA*, and *rps1* are regulated by LexA. Absence of AnLexA-Box upstream of the *fus-tufA* operon (Table 2) accounts for the two genes not being regulated by LexA. While in case of *prk* and *alr0803*, the predicted AnLexA-Box is distant (over 100 bases) from the predicted promoter (Table 2) and thus is unlikely to be regulated by LexA, even though the LexA is capable of binding to it as shown by EMSA studies (Fig. 3). The discrepancies observed between changes at transcript and protein level is likely to be due to significant difference in size of the protein spots and the actual molecular mass of these proteins, (Table 1) with the actual intact protein not having been detected/identified.

**Fig. 4.**
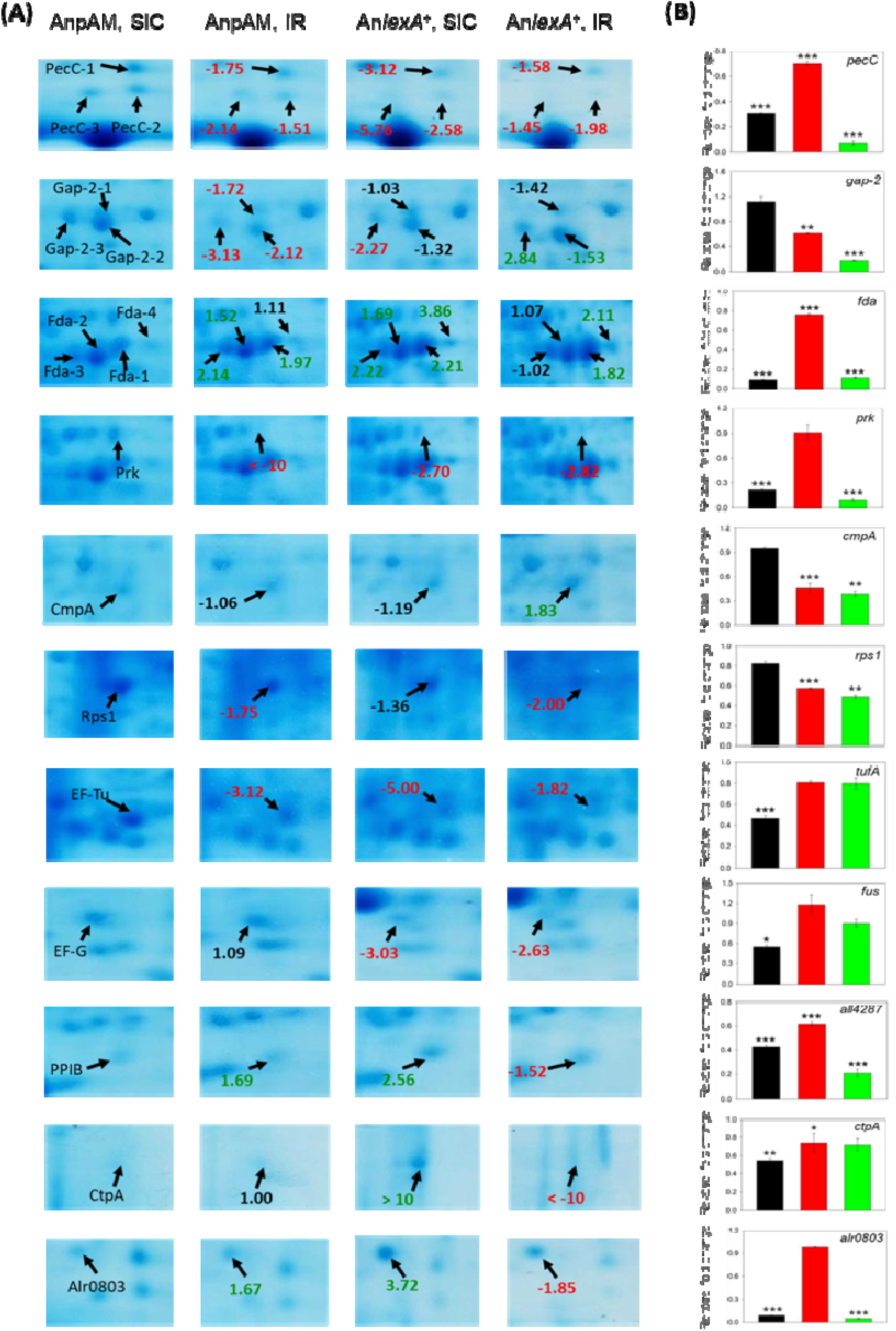
Comparison of (A) proteomic and (B) transcript level changes for select proteins and corresponding genes as indicated. (A) The protein spots are enlarged images of the corresponding region in Fig. 1, with the names indicated in the panel under AnpAM, SIC, while fold-change as indicated in Table 1 are shown in the panels corresponding to AnpAM, IR; An*lexA*^+^, SIC; and An*lexA*^+^, IR. (B) The change in transcript levels determined through quantitative reverse transcriptase (qRT)-PCR of the corresponding genes (i) upon IR in AnpAM cells with respect to AnpAM, SIC is shown black, (ii) upon LexA overexpression in An*lexA*^+^, SIC with respect to AnpAM, SIC in red, and (iii) upon IR in An*lexA*^+^ cells with respect to An*lexA*^+^, SIC in green. The cut-offs of a fold change of 1.3 and *p* < 0.05 was applied to filter expressed genes with significant differences and high confidence (differentially expressed genes are indicated with asterisks). Based on Student’s t-test with Tukey’s post-hoc comparison test, *, ** and *** correspond to *p* < 0.05, *p* < 0.01, and *p* < 0.001, respectively.

A good agreement was observed between change in abundance of protein and that of the transcript upon γ-irradiation both in AnpAM and An*lexA*^+^ cells for *pecC*, *prk*, *rps1*, and *ctpA* for *tufA* (EF-Tu) and *fus* (EF-G) in AnpAM and for *gap-2*, *all4287*, and *alr0803* in An*lexA*^+^ cells (Fig. 4). Upon irradiation, the *cmpA* transcript levels were enhanced in AnpAM but lowered in An*lexA*^+^, which could be due to the inhibition of transcription of *cmpA* by LexA as observed in control cultures of An*lexA*^+^ (Fig. 4).

### 3.3 **γ**-radiation and LexA-induced changes in photosynthetic parameters and photosynthetic complex stoichiometry in Anabaena

RLCs of variable chlorophyll *a* fluorescence corresponding to PSII revealed higher quantum yield of PSII [Y(II)] in An*lexA*^+^ compared to AnpAM under both SIC and IR conditions, the values showing similar decrease in IR in both the cultures (Fig. 5A). No significant change was observed for photochemical quenching coefficients, qP and qL between the different cultures used (Fig. 5B, C). ETR(II) was found to be similar for AnpAM and An*lexA*^+^, SIC cultures under low irradiance, but was higher in An*lexA*^+^ at high light irradiance, and decreased upon γ-radiation for both the cultures, the magnitude of decrease being higher for AnpAM compared to An*lexA*^+^ (Fig. 5D). The unregulated heat-dissipation [Y(NO)] was similar for AnpAM and An*lexA*^+^ cultures and increased upon γ-radiation (Fig. 5E).

**Fig. 5.**
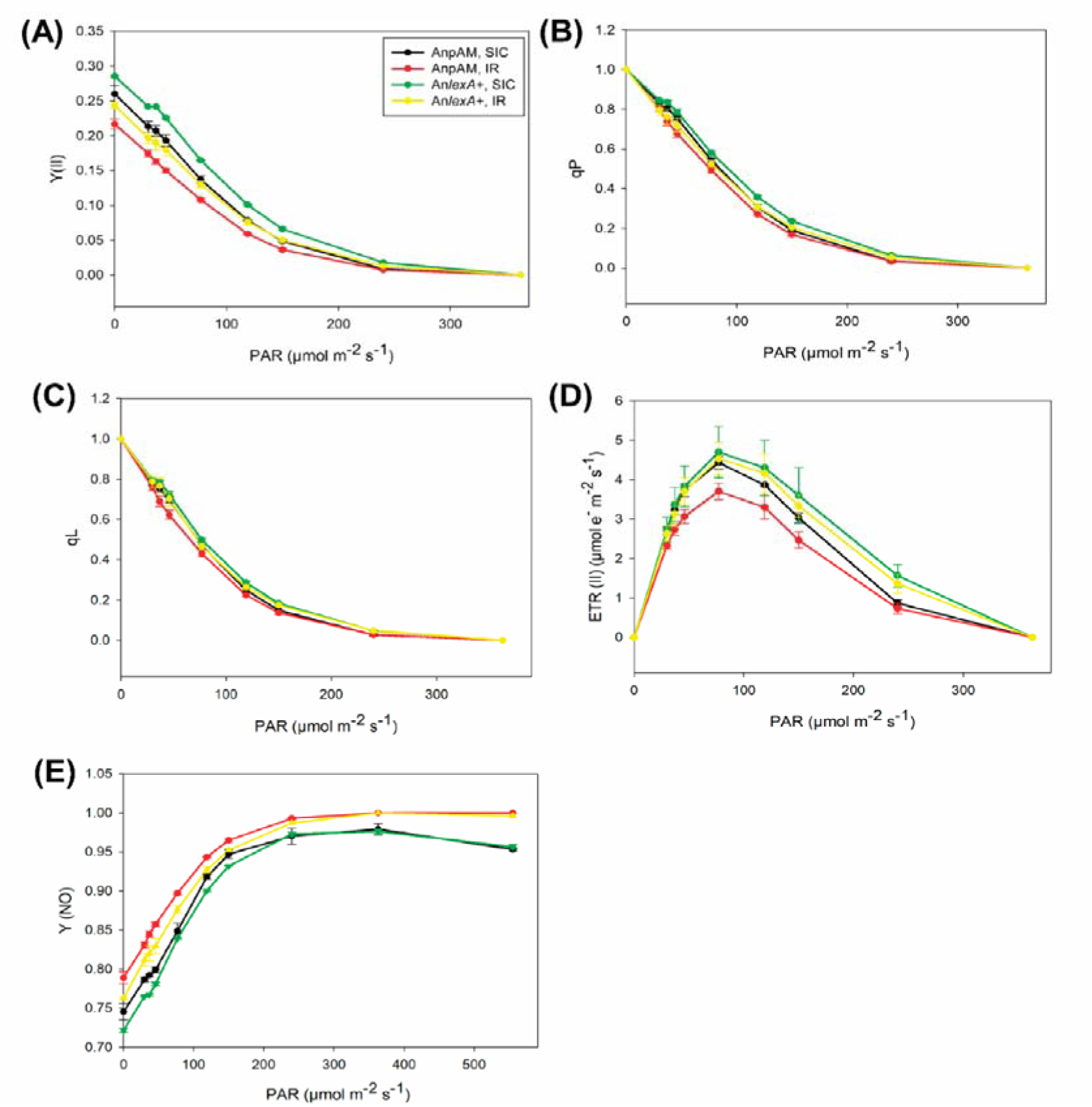
Rapid light curves (RLCs) of AnpAM and An*lexA*^+^ cells corresponding to sham-irradiated controls (SIC) and 6 kGy γ-irradiation (IR) were evaluated for the following photosynthetic parameters of PSII (A) Y(II) [effective quantum yield of photochemical energy conversion], (B and C) coefficient of photochemical quenching based on (B) puddle model (qP) and (C) lake model (qL), (D) ETR(II) [electron transfer rate], and (E) Y(NO) [quantum yield of non-regulated energy dissipation]. Each value represents the mean ± SE of at least three biological replicates (n = 3).

The identification of the 10 bands corresponding to the five main photosynthetic complexes namely, photosystem I (PSI), photosystem II (PSII), NAD(P)H dehydrogenase-1 (NDH-1), cytochrome *b*_6_*f* (Cyt *b*_6_*f*), and ATP synthase on the BN-PAGE gel (Fig. 6) were done based on earlier published work (Herranen et al., 2004; Gao et al., 2016; Srivastava et al., 2021a,b). In SIC, overexpression of LexA resulted in a visible decrease in the intensities of PSI supercomplexes, tetramer, and Cyt *b_6_f* monomer (Fig. 6), the fold-decrease being 1.59, 1.38, and 1.33, respectively (Supplementary Table S4), and increase in the levels of PSI and PSII monomers, and Cyt *b_6_f* dimer by 1.75-, 1.31-, and 1.49-fold, respectively (Fig. 6, Supplementary Table S4). Exposure to γ-radiation resulted in increase in the levels of CP43-lesss PSII complex by 1.57- in AnpAM while decrease in its level in An*lexA*^+^ by 1.31-fold, and decrease in level of PSI super-complexes and PSI tetramer by 1.38- and 1.41-fold, respectively in An*lexA*^+^ cells (Fig. 6, Supplementary Table S4).

**Fig. 6.**
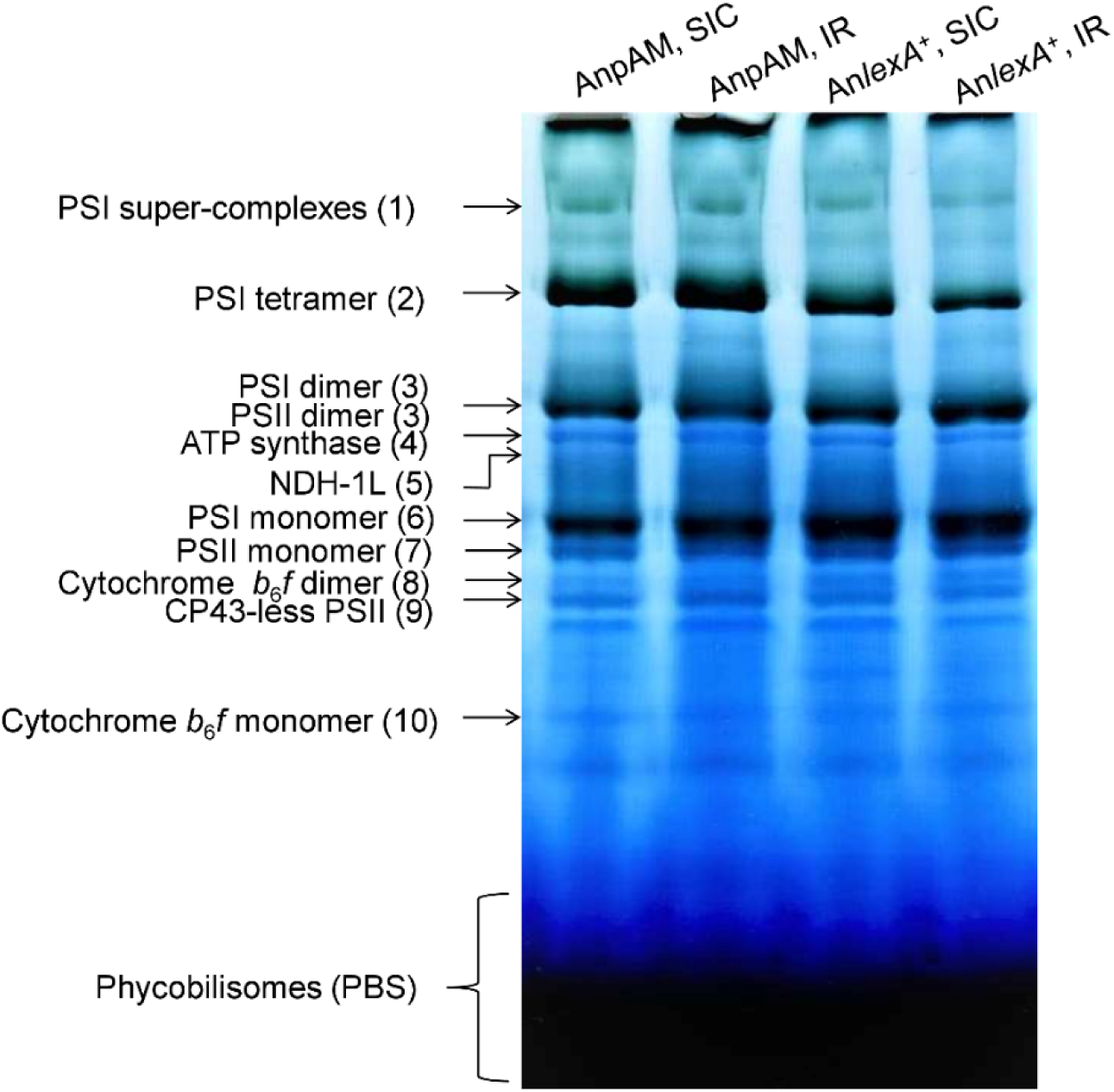
Thylakoid membrane proteome profiling of sham-irradiated control (SIC) and γ-irradiated (IR) cultures of recombinant *Anabaena* strains AnpAM and An*lexA*^+^ using blue native (BN-PAGE). The different oligomeric forms of the photosynthetic complexes are labeled and also numbered. Quantification of band signal intensities using ImageJ software (http://rsb.info.nih.gov/ij) for all 10 bands is given in Supplementary Table S4.

### 3.4 Comparison of the proteome of AnpAM and AnlexA^+^ cells as function of short-term exposure to dark conditions

In the present study, the sham irradiated controls were kept in dark to mimic the conditions of irradiated cultures, while in our earlier study (Kumar et al., 2018), both the cultures i.e., AnpAM and An*lexA*^+^ were kept under illuminated conditions prior to proteomic analysis. In order to assess the contribution of dark conditions (∼1.6 h), we compared the fold change of spots identified in the current study in response to LexA overexpression (Fig. 1A,C), with that observed earlier (Kumar et al., 2018), which is tabulated in Supplementary Table S5. Of the total 41 spots identified during the present study, 21 spots showed similar trends upon comparing control AnpAM and An*lexA*^+^ cells kept in dark for ∼1.6 h (Fig.1A, C) and those which were under constant illumination (Kumar et al., 2018), though the fold change was different (Supplementary Table S5). Of the remaining 20 protein spots, 9 were not detected in either AnpAM or An*lexA*^+^ gels during the earlier experiment (Kumar et al., 2018), and these corresponded to three protein spots of CpcC, one each of Fld, Rps8, PPIB, PRK, CtpA, and Alr0882 (Supplementary Table S5). Opposite trend was observed for one protein spot of Fda and RpoC2 (Supplementary Table S5). This indicated that light does influence the extent of regulation by LexA, and this could be due to the expression of the LexA from the light-inducible promoter, P*_psbA1_*, which would not be active while the cells were maintained in dark during the present experiment.

## 4. Discussion

Proteins are the key executors of cellular processes and are essential for maintaining cellular homeostasis (Liu et al., 2019). Every organism reprograms its cellular machinery at the protein level to adapt to challenging conditions. This would also be true for cyanobacteria, the oldest oxygen-producing photosynthetic organisms on Earth dating back at least 2.6 billion years (Hedges et al., 2001), which would have adapted this approach to survive high radiation conditions prevailing then. The cyanobacterium, *Anabaena* achieved high radioresistance by protecting its physiological functions by undergoing proteomic changes (Singh and Apte, 2018) and an efficient DNA repair (Singh et al., 2013). While several of the DNA repair genes have been speculated or proven to be regulated by LexA owing to the presence of AnLexA-Box (Kirti et al., 2017; Kumar et al., 2018; Pandey et al., 2018; Rajaram et al., 2020), it remained to be seen if LexA could also regulate some of the genes corresponding to the proteomic changes observed, and thus function as one of the global regulators for radiation response in *Anabaena*. The ability of LexA to modulate radiation and oxidative stress tolerance (Kumar et al., 2018) led credence to its possible involvement in regulating genes involved in tolerance to these stresses.

### 4.1 Modulation of photosynthetic responses of Anabaena under **γ**-radiation stress as a function of LexA levels

Proteomic analysis of *Anabaena* to γ-radiation in the absence or presence of high levels of constitutively expressed LexA revealed that over 35% of the proteins showing change in abundance under these conditions, were from the functional category of photosynthesis and C-metabolism (Table 1, Fig. 2C). Of the genes coding for photosynthetic pigments, down-regulation of *cpcB* operon and up-regulation of *apcE* by LexA has been demonstrated earlier (Kumar et al., 2018). The *pecC* gene which is present as an operon with downstream *pecE* gene was found to be down-regulated at transcript and protein level both upon γ-irradiation and LexA overexpression in *Anabaena* (Fig. 4), with the down-regulation by LexA being through the binding of LexA to the AnLexA-box (Fig. 3). Two AnLexA-boxes were detected upstream of the *pecC* start codon, one box overlapping with the predicted promoter region and the other immediately downstream of it (Table 2) suggestive of a tight regulation by LexA. The PSII photophysiology dynamics of the recombinant *Anabaena* strains, AnpAM and An*lexA*^+^ under IR conditions revealed similar rates of decrease in PSII photochemical efficiency, fraction of open reaction centres (RCs), and electron transport and increase in non-regulated energy dissipation owing to closed PSII RCs (Fig. 5). However, under control (sham-irradiated) conditions, An*lexA*^+^ exhibited greater PSII photochemical efficiency and lower energy dissipation than AnpAM (Fig. 5). The observed photophysiological changes reflected on the stoichiometry dynamics of photosynthetic complexes in AnpAM and An*lexA*^+^ under γ-radiation (Fig. 6). Of the 10 identified oligomeric forms of five photosynthetic complexes, the changes observed upon γ-irradiation was similar for 7 complexes in case of AnpAM and An*lexA*^+^ cells, with distinct differences observed for PSI super-complexes (Band 1) and PSI tetramer (Band 2), as well as CP43-less PSII (Band 9), which showed a decrease in An*lexA*^+^ but increase in AnpAM cells upon irradiation (Fig. 6, Supplementary Table S4). Of these, the CP43-less PSII is an intermediary in both the PSII repair cycle and de novo PSII assembly (Shi et al., 2021), and its accumulation is indicative of increased PSII photodamage, reduced PSII efficiency, and hence decreased linear electron flow (LEF) (Theis and Schroda, 2016). Thus, the lower accumulation of CP43-less PSII in irradiated An*lexA*^+^ cells compared to AnpAM cells could explain the higher PSII photochemical efficiency of An*lexA*^+^ (Fig. 5A). The observed changes in the stoichiometry of these complexes in sham-irradiated controls upon LexA overexpression (Fig. 6, Supplementary Table S4) was similar to that observed for normal controls (Srivastava et al., 2022). Under stress conditions, when LEF along PSII is inhibited, cyanobacteria induce NDH-dependent cyclic electron flow (CEF) around PSI to offer photo-protection (Gao et al., 2016; Srivastava et al., 2021a). Lower levels of PSI tetramer under IR in *AnlexA*^+^ compared to AnpAM (Fig. 6, Supplementary Table S4) suggests higher photodamage to PSI, due to higher flow of PSII-trapped electrons towards PSI, which is also an indicative of the inability of An*lexA*^+^ to restrict LEF and induce NDH-mediated CEF. The lower accumulation of NDH-PSI super-complex in An*lexA*^+^ than in AnpAM under both SIC and IR conditions (Fig. 6, Supplementary Table S4) was again an indicative of slower LEF restriction and decreased NDH-mediated CEF around PSI in An*lexA*^+^. Oxygenic photosynthetic organisms decrease PSII quantum efficiency and restrict electron transport in ETC downstream of the PSII in response to abiotic stress conditions, including γ-radiation, to maintain coordination between light and dark photosynthetic reactions (Choi et al., 2021; Srivastava et al., 2021a), as the enzymes involved in Calvin-Benson-Bassham (CBB) cycle are very sensitive to oxidative stress (Krupa et al., 1993; Muhammad et al., 2021). Thus, the inability to inhibit PSII photochemical efficiency sufficiently and inflicting higher damage to PSI during γ-radiation in An*lexA*^+^ cells, contributes to the lower radiation tolerance of An*lexA*^+^ compared to AnpAM.

### 4.2 LexA as one of the regulators of **γ**-radiation stress response in Anabaena

Cytosolic proteome reprogramming of AnpAM and An*lexA*^+^ under IR conditions revealed the differential accumulation (>1.5-fold, *p* < 0. 05) of 41 DAP spots comprising 29 distinct proteins, which were divided into six FCs (Figs. 1, 2, 4, Table 1, Supplementary Table S3). Except for *fus* and *tufA* encoding for EF-G and EF-Tu, respectively and comprising a single operon, AnLexA-Box was found upstream of all other identified genes, including three genes *cpcC*, *rps8*, and *all2133* which are transcribed from promoter of their upstream genes (Table 2). This indicated that LexA could be one of the regulators of radiation-responsive genes in *Anabaena*.

#### 4.2.1 Photosynthetic and C-metabolism genes

Of the 12 DAPs corresponding to 5 proteins (Table 1), CpcB and ApcE are subunits of chromophore-associated water soluble phycobiliproteins (PBPs). In contrast, CpcC, CpcG4, and PecC are nonchromophoric and hydrophobic linker polypeptides, required to organize PBPs into supramolecular complexes known as phycobilisome (PBS), which are coated on the outer surface of TMs in form of regular arrays (Bryant et al., 1979; Singh et al., 2015). Of these *apcE* has been shown to be up-regulated by LexA in *Anabaena* earlier (Kumar et al., 2018), while CpcG4 showed higher abundance in An*lexA*^+^ cells (Fig. 1, Table 1) and with the presence of AnLexA-box overlapping with the predicted -10 region (Table 2), the *cpcG4* gene is predicted to be regulated by LexA. On the other hand, *cpcB*, *cpcC*, and *pecC* which show decreased accumulation in An*lexA*^+^ cells (Figs. 1, 4) have been shown to be down-regulated by LexA in *Anabaena* at transcriptional level also (Fig. 4, Kumar et al., 2018). Such varied regulation of the PBS component genes by LexA could either result in change in stoichiometry of components of PBS or decreased levels of PBS complex based on lower availability of proteins whose expression is down-regulated by LexA. Exposure to γ-radiation stress resulted in decreased abundance of all these proteins (Table 1), which could be a means of protecting its photosystem, since, the declination of PBPs and further decoupling of PBS have been suggested to provide photo-protection by decreasing the transfer of absorbed light energy to the PSII RCs (Tamary et al., 2012). Another photosynthesis-associated protein identified was the electron transport protein Flavodoxin (Fld), which showed increased accumulation in the presence of LexA, as well as in response to γ-radiation (Fig. 1, Table 1). The corresponding gene (*alr2405*) possessed multiple AnLexA-Boxes near its predicted -10 promoter (Table 2), and is thus expected to be regulated by LexA. Of the several Fe-containing photosynthetic proteins, *petH* and *all2919* (Fd) are negatively regulated by LexA in *Anabaena* (Kumar et al., 2018; Srivastava et al., 2022), thus the enhanced expression of Fld in the presence of LexA could aid the cells in electron transport reactions requiring Fe-containing proteins. Fld was initially discovered in cyanobacteria growing under iron-limiting conditions (Smillie et al., 1965), and has been shown to be induced and substitute for ferredoxin (Fd) under environmental stress and iron deprivation conditions (Lodeyro et al., 2012). Thus, the increased levels of Fld under IR conditions could be a means of mitigating the stress. In An*lexA*^+^ cells, decreased expression of *galE* and *talB* (Kumar et al., 2018), *gap-2* (Fig. 4) has been observed at transcript and protein level, while for *fda*, the transcript level decreases, while the total protein spots corresponding to Fda are higher in abundance in the cells, suggesting higher cellular stability of the Fda protein. This along with the observed increased abundance of Rpe (Fig. 1, Table 1) would allow functioning of the phosphopentose pathway (PPP) as a means of NADP/ATP pool management with glycolysis being lower due to lower abundance of some of the key proteins in the pathway. The observed proteomic changes in the FCI proteins in AnpAM cells upon γ-irradiation (Figs. 1, 2, 4, Table 1) largely agrees with the proteomic changes observed in the wild type *Anabaena* cells earlier (Singh and Apte, 2018). The profile was similar for IR-treated An*lexA*^+^ cells, as well except for *cmpA* whose transcription was repressed in An*lexA*^+^ cells under control as well as IR conditions though the degraded protein spot of ∼39 kDa, as against the actual mass of ∼50 kDa, showed higher accumulation in An*lexA*^+^, IR cells (Figs. 1, 4). CmpA is a bicarbonate-binding protein and a key component of CO_2_-concentrating mechanism (CCM) (Koropatkin et al., 2007), and could be playing a role in maintaining C-balance in An*lexA*^+^ cells. The observed regulation of photosynthetic genes by LexA, both as an activator and a repressor for different genes involved in photosynthesis is in agreement with our earlier findings of the global regulatory role of LexA in *Anabaena*, both as an activator and a repressor (Kumar et al., 2018). However, we are yet to identify the distinguishing features of the promoter element that allow the binding of LexA to AnLexA-Box to differentiate its role as an activator or repressor.

#### 4.2.2 Oxidative stress alleviation genes

An*lexA*^+^ cells have been earlier shown to be sensitive to oxidative stress (Kumar et al., 2018), but as a regulatory protein, LexA maintains a fine balance in terms of regulating the expression of oxidative stress alleviating proteins. The decreased abundance of PrxA and CBS (Alr4416) (Fig. 1, Table 1), of which *prxA* was earlier shown to be down-regulated by LexA along with two other genes, *sodA*, *katB* on one hand, and the increased abundance of Trx2 (Fig. 1, Table 1) along with that of ApcE (Kumar et al., 2018) on the other hand, allows An*lexA*^+^ cells in maintaining a fine balance of ROS to allow it to survive under normal growth conditions. In response to radiation stress, both AnpAM and An*lexA*^+^ cells respond by enhancing the levels of CBS in addition to Trx2, while PrxA levels decrease (Fig. 1, Table 1). The antioxidant defense proteins/enzymes Trx2, PrxA, and CBS have been shown to increase under a variety of abiotic stresses and protect essential cellular components from ROS-induced oxidation (Mishra et al., 2009; Srivastava et al., 2021b). Another important protein shown to have decreased abundance in An*lexA*^+^ cells is the stress-inducible DNA-binding protein DpsA, which is known to protect DNA integrity and cellular viability during environmental stress and nutritional deficiency (Karas et al., 2015), and its deletion mutation in *Synechococcus* sp. PCC7942 resulted in them being highly susceptible to all photooxidative stress conditions, including high light and paraquat treatment (Dwivedi et al., 1997). The AnLexA-Box was found to be present immediately upstream of the predicted -10 region of *dpsA* (Table 2), and is thus likely to be regulated by LexA. Lower DpsA levels in An*lexA*^+^ may affect its stress tolerance to different stresses, as observed earlier (Kumar et al., 2018).

#### 4.2.3 Transcription, translation, and protein folding genes

Of the identified proteins involved in mRNA and protein synthesis and modifications (FCII and FCIV), all the corresponding genes except for *fusA-tufA* operon coding for EF-G and EF-Tu, respectively possessed AnLexA-Box in the vicinity of the -10 region (Table 2), indicating that LexA would regulate them. The absence of regulation of the two genes of this operon by LexA was confirmed by transcript analysis, where in no significant change was observed in their levels in An*lexA*^+^ cells compared to AnpAM under control conditions (Fig. 4). Both the genes were, however, found to be down-regulated at the transcriptional level in response to γ-radiation (Fig. 4), conforming to the earlier observed protein data in wild type cells (Singh and Apte, 2018). RpoC2, which is involved in gene transcription, exhibited lower abundance in response to γ-radiation as well in An*lexA*^+^ cells (Fig. 1), presence of AnLexA-Box overlapping with the -10 region (Table 2) confirming its regulation by LexA. This could contribute to observed lower transcript yield in *Anabaena* cells exposed to γ-radiation as well as in An*lexA*^+^ compared to AnpAM (data not shown). While the elongation factors involved in protein synthesis were not found to be regulated by LexA, the genes coding for ribosomal proteins were, as indicated by the presence of AnLexA-Box upstream of *rps1* and *rpl8-rps5* genes/operon in the vicinity of the -10 region (Table 2) and the observed down-regulation of *rps1* transcription (Fig. 4). Both the proteins were found to be down-regulated upon γ-radiation (Fig. 1) in tune with the observed decreased protein synthesis as a response to IR. The chaperone, PPIB which catalyzes the cis-trans isomerization of proline imidic peptide bonds in oligopeptides and facilitates protein folding (Hassidim et al., 1992), was found to increase in abundance upon LexA overexpression and in response to IR in AnpAM cells but decreased in IR-stressed An*lexA*^+^ (Figs. 1, 4). However, the transcription of this gene was found to be down-regulated by both IR and LexA overexpression (Fig. 3), suggesting higher stability of the protein in the cells. The down-regulation of *all4287*, coding for PPIB by LexA (Fig. 4) was through the binding of the LexA protein to the AnLexA-Box (Fig. 3). Reduced transcription of *all4287* by IR, as well as LexA (Fig. 4) could be responsible for the lower abundance of PPIB in An*lexA*^+^ upon IR (Figs. 1,4). Degraded form of carboxyl-terminal protease, CtpA could be detected only in An*lexA*^+^ cells under control conditions (Figs. 1,4), but the transcript was found to be down-regulated by both IR and LexA, through its AnLexA-box (Table 2), and IR (Fig. 4). CtpA is needed to convert the precursor PSII RC protein D1 to mature D1 by eliminating the short C-terminal extension, and keeping PSII active (Anbudurai et al., 1994). The loss of D1 maturation by CtpA has also been demonstrated to impair the assembly of PSII core proteins D2, CP43, and CP47, with the incorporation of CP43 being the most impacted (Shi et al., 2021). Thus, it could account for the observed photodamage of PSII in response to IR.

#### 4.2.4 Fatty acid synthesis genes

The abundance of ENR (*fabI*, *all4391*) and ACS (*all4257*) was found to be enhanced upon overexpression of LexA but was unaffected by IR stress (Fig. 1, Table 1), and AnLexA-Box was detected in the vicinity of -10 region of both the genes (Table 2), suggesting their regulation by LexA. ENR or FabI has been shown to play a determining role in the final steps of elongation of fatty acid biosynthesis (Heath and Rock, 1995), while ACS is involved in the generation of acetyl CoA from acetate, which is used in several metabolic processes including fatty acid biosynthesis. In the unicellular cyanobacterium *Synechocystis* sp. PCC6803, the LexA was found to down-regulate the expression through binding to upstream region of *fabD*, *fabH*, and *fabF* which are involved in the initiation steps of fatty acid biosynthesis and *fabG* involved in the first reductive step during elongation, and bound to the upstream region of *fabI* and *fabZ* (Kizawa et al., 2017). Presence of AnLexA-Box upstream of the *fabH*, *fabH*, *fabF*, and *fab*Z of *Anabaena* has been reported earlier (Kumar et al., 2018), though its regulation has not been demonstrated. Thus, the LexA could play an important regulatory role in fatty acid biosynthesis in cyanobacteria.

#### 4.2.5 Others

A few other important proteins were found to exhibit enhanced or decreased accumulation in response to LexA overexpression and IR stress or both. One of them is a putative PCD coded by *asr4549*, which is involved in recycling of the pterin cofactors, which are generated by aromatic amino acid hydroxylases (Naponelli et al., 2008). PCD has also been shown to be involved as a RuBisCo assembly factor in the α-carboxysomes of *Thiomonia intermedia*, a chemoautotroph, and homologs of the α-carboxysomes have been found in other bacteria, alga and higher plants (Wheatley et al., 2014). The protein was found to show enhanced abundance upon overexpression of LexA and in response to γ-radiation (Fig. 1, Table 1) and could enhance the formation of carboxysomes and protect RuBisco, an important protein for the photosynthetic bacterium. Four other proteins identified are annotated as either hypothetical or unknown proteins in the database. Of these, All2133 exhibited high similarity with PII-like signaling protein, SbtB which is a part of CCM (Wang et al., 2004). The higher accumulation of All2133 level in An*lexA*^+^ compared to AnpAM under both control and IR conditions (Fig. 1, Table 1), could be contributing to the higher C-assimilation in An*lexA*^+^ cells. Its regulation by LexA was confirmed bioinformatically through the presence of AnLexA-Box in the vicinity of the -10 region (Table 2). Alr0803 was similar to signal transduction histidine kinase and has been earlier shown to be regulated in response to abiotic stresses such as salt, UV-B radiation, heavy metal, desiccation (Rai et al., 2013; Singh et al., 2015; Sen et al., 2017; Babele et al., 2019). Though the protein spots corresponding to Alr0803 showed increased abundance upon IR in AnpAM cells, the size of the identified protein was much smaller and may be only the increase in the degraded product. On the other hand, RT-PCR analysis, confirmed down-regulation of the gene in response to IR in both AnpAM and An*lexA*^+^ cells (Fig. 4). The difference in the spot size and actual protein size may also account for the observed increase in abundance on protein gels (Figs.1, 4), but no change in transcript levels (Fig. 4) in An*lexA*^+^ cells compared to AnpAM under control conditions. The upstream region of *alr0803* possessed AnLexA-Box (Table 2) confirmed by the binding of LexA to the corresponding DNA fragment (Fig. 3). However, the distance between the AnLexA-Box and predicted -10 region is over 100 bases (Table 2), and thus, may not facilitate regulation by LexA. Thus, *alr0803* could be a universal stress-responsive gene responding to γ-radiation stress as well. Alr1144 has not been found to be similar to any of the known proteins. The presence of AnLexA-Box upstream of *alr1144* in the vicinity of the -10 region (Table 2) accounts for its regulation in An*lexA*^+^ compared to AnpAM under unstressed conditions (Fig. 1, Table 1). Alr0882 showed increased abundance in response to IR and LexA overexpression in *Anabaena* (Fig. 1, Table 1), and is likely to be regulated by LexA based on the presence of AnLexA-Box near to the -10 region (Table 2). Alr0882 showed similarity with universal stress protein (USP) domain-containing proteins, which are known to be up-regulated in response to any abiotic stress. Thus, many of the proteins regulated in response to irradiation also exhibited regulation in response to LexA overexpression indicating that LexA could be one of the regulators of IR-responsive genes in *Anabaena*.

## Conclusions

To cope with γ-radiation stress, cyanobacteria temporarily reduce photosynthesis and growth while activating defence, detoxification, and repair mechanisms; and such changes need to be transcriptionally controlled. Since, the repertoire of genes undergoing changes in expression in response to γ-radiation is large; several regulators may be involved in this process, working either individually or in tandem. However, not much information is available on the regulators of IR response in the radioresistant cyanobacterium, *Anabaena* sp. PCC7120. The earlier predicted global regulatory role of LexA in *Anabaena* (Kumar et al., 2018) was evident in response to γ-radiation as well, as it was found to regulate several genes corresponding to proteins whose abundance was affected in response to IR. The 41 differentially abundant spots (≥ 1.5-fold change) in control and irradiated cultures of AnpAM and An*lexA*^+^ corresponded to 29 proteins, which included 13 new proteins compared to that identified earlier for *Anabaena* in response to γ-radiation (Singh and Apte, 2018), 22 new proteins regulated by LexA under control growth conditions compared to those identified earlier in An*lexA*^+^ (Kumar et al., 2018). The study revealed LexA-mediated regulation of genes regulating photosynthesis directly (Srivastava et al., 2022) or indirectly through stabilising and protecting the photosynthetic complexes, and those involved in C-catabolism. LexA also regulated oxidative stress-responsive genes differentially to maintain a fine balance in dismutating or removing ROS. While this fine balance of ROS and C-status is maintained to the advantage of the cells under normal growth conditions, upon overexpression of LexA, exposure to IR disrupts this balance resulting in lowering of radiation tolerance of An*lexA*^+^ cells (Kumar et al., 2018), along with the predicted down-regulation of genes involved in DNA repair by LexA (Rajaram et al., 2020). A schematic representation of the regulation of different genes by LexA in response to γ-radiation stress is shown in Fig. 7. Identification of LexA as one of the regulators is the first step in identifying the regulatory network and elements contributing to radiation-response of *Anabaena* through modification of its transcriptome and proteome. And it could thus initiate further research in this direction.

**Fig. 7.**
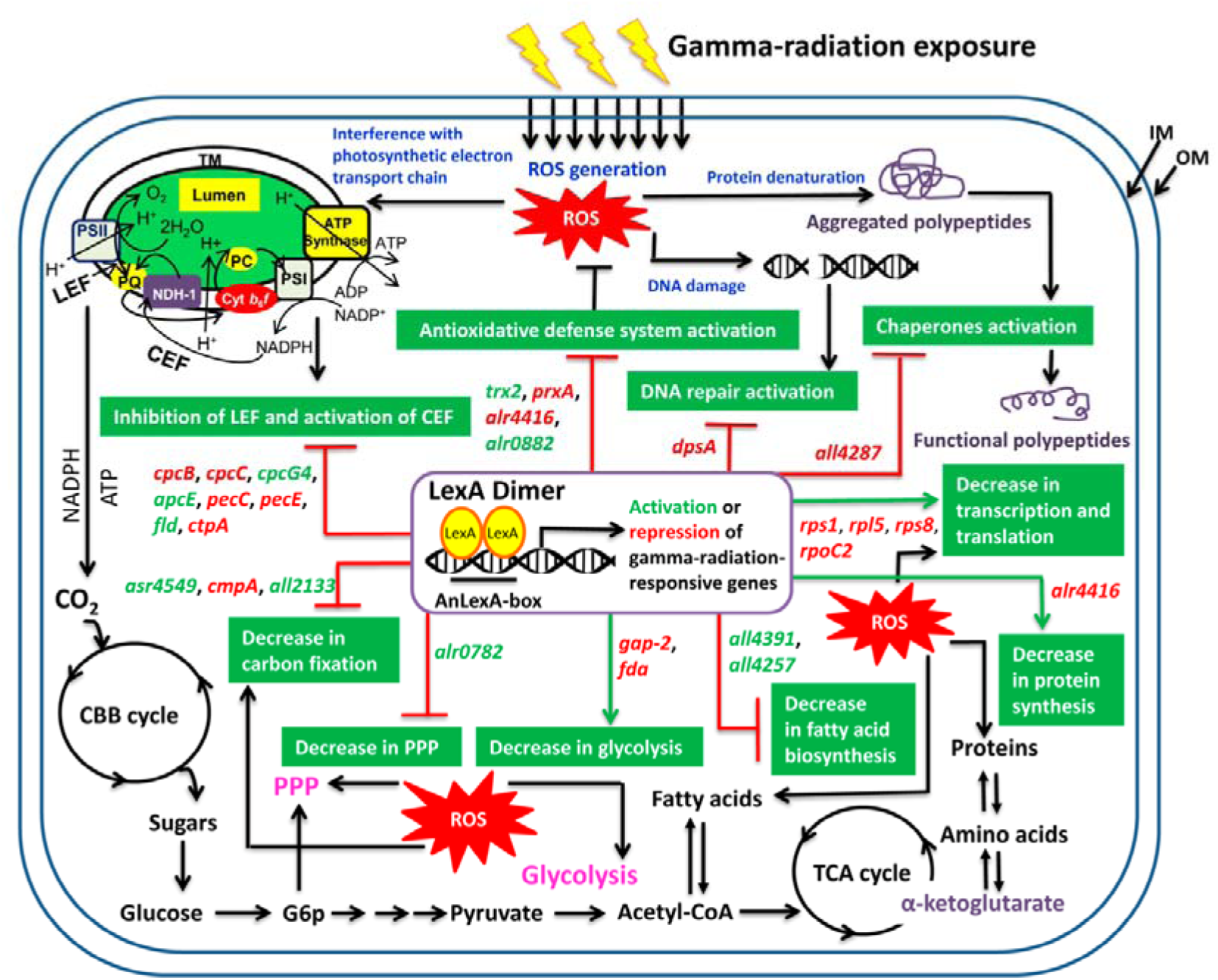
A schematic representation of LexA regulation on different γ-radiation-responsive genes, as well as radiation stress response of *Anabaena* sp. PCC7120. LexA-activated or repressed genes (gene symbols are italicized) in response to γ-radiation stress are highlighted in green or red, respectively. The proteins encoded by these genes are listed in Table 1. The adverse effects of the γ-radiation in *Anabaena* are written in blue, whereas adaptive responses induced to cope with radiation stress are written in white in green boxes as per the findings of this study, as well as Singh et al. (2013) and Singh and Apte, (2018). The possible regulation of LexA on γ-radiation stress response of *Anabaena* through activation or repression of different radiation-responsive genes is illustrated by a green arrow (→) for induced responses and a red inhibitory arrow (T) for repressed responses. (ATP, adenosine triphosphate; CBB, Calvin-Benson-Bassham; CEF, cyclic electron flow; Cyt *b*_6_*f*, cytochrome *b*_6_*f* complex; G6p, glucose-6-phosphate; IM, inner membrane; LEF, linear electron flow; NADP^+^, nicotinamide adenine dinucleotide phosphate; NADPH, nicotinamide adenine dinucleotide phosphate hydrogen; NDH, NAD(P)H dehydrogenase; OM, outer membrane; PC, plastocyanin; PPP, phosphopentose pathway; PSI, photosystem I; PSII, photosystem II; ROS, reactive oxygen species; TCA cycle, tricarboxylic acid cycle; TM, thylakoid membrane).

## Acknowledgments

A.S. thanks to Department of Science and Technology-Innovation of Science Pursuit for Inspire Research (DST–INSPIRE, New Delhi, India) fellowship and S. B. thanks to Council of Scientific & Industrial Research (CSIR, New Delhi, India) fellowship for Ph.D. Y.M. acknowledges the financial support provided by the Department of Atomic Energy-Board of Research in Nuclear Sciences (DAE-BRNS, Mumbai, India) (Grant number: 37(1)/14/12/2017-BRNS/37209. He also acknowledges Institute of eminence (IOE) incentive grant, Banaras Hindu University (BHU, Varanasi, India) (R/Dev/D/IOE/Incentive/2021-22/32401). We wish to thank Dr. Rachna Agarwal (Molecular Biology Division-Bhabha Atomic Research Centre, Mumbai, India) for providing Dual-PAM fluorimeter (DUAL-PAM100, GmbH Germany) facility for PSII photophysiological measurements. For statistical analysis, we are thankful to Prof. Ram Sagar (Department of Botany, Banaras Hindu University, Varanasi, India). We would also like to express our gratitude to the Head and the Programme Coordinator (CAS) in Botany and Interdisciplinary School of Life Science (ISLS), at Banaras Hindu University, Varanasi, India, for providing other instrumental support.

## Author Statement

A.S., A.K., H.R. and Y.M. conceived the idea and designed the experiments. A.S., A.K., and S.B. conducted the experiments. V.S. did the mass spectrometry analysis. A.S., V.S., A.K., H.R., and Y.M. analyzed the data and wrote the manuscript with help of remaining authors.

## Supplementary files

**Table S1** List of primers used for transcript analysis

**Table S2** List of primers used for promoter analysis (EMSA)

**Table S3** List of spots showing changes in abundance in recombinant *Anabaena* strains in response to IR or LexA overexpression

**Table S4** Quantification of band signal intensities in BN-PAGE gel using ImageJ software (http://rsb.info.nih.gov/ij) based on densitometry analysis

**Table S5** Comparative analysis of fold change of protein spots in the current study with the corresponding spots in the data published earlier by Kumar et al. (2018)

## Abbreviations

ATP synthase: Adenosine triphosphate synthase
BN-PAGE: Blue native-polyacrylamide gel electrophoresis
CBB: Calvin-Benson-Bassham
CCM: CO_2_-concentrating mechanism
cDNA: Complementary DNA
CEF: Cyclic electron flow
Cyt *b*_6_*f*: Cytochrome *b*_6_*f* complex
DAP: Differentially accumulated protein
EMSA: Electrophoretic mobility shift assay
ETR(II): Electron transport rate of PSII
FCs: Functional categories
IEF-SDS PAGE: Isoelectric focusing-sodium dodecyl sulphate-polyacrylamide gel electrophoresis
IR: γ-irradiated
kGy: kiloGrays
LEF: Linear electron flow
NDH: NAD(P)H dehydrogenase
PAR: Photosynthetically active radiation
PBPs: Phycobiliproteins
PBS: Phycobilisome
PCA: Principal component analysis
pI: Isoelectric point
PPFD: Photosynthetic photon flux density
PPP: Phosphopentose pathway
PSI: Photosystem I
PSII: Photosystem II
qL: Lake model-based coefficient of photochemical quenching
qP: Puddle model-based coefficient of photochemical quenching
qRT-PCR: Quantitative reverse transcriptase-Polymerase chain reaction
RCs: Reaction centres
RLCs: Rapid light curves
ROS: Reactive oxygen species
RuBisco: Ribulose-1,5-bisphosphate carboxylase/oxygenase
SE: Standard error
SIC: Sham-irradiated control
TM: Thylakoid membrane
USP: Universal stress protein
Y(II): Effective quantum yield of photochemical energy conversion in PSII
Y(NO): Quantum yield of non-regulated energy dissipation in PSII

