## Supplementary files for "Gamma (γ)-radiation stress response of the cyanobacterium *Anabaena* sp. PCC7120: Regulatory role of LexA and photophysiological changes"

#: Equal First Authors

**Table S1** List of primers used for transcript analysis

| **S. N.** | **Genes**  **encoding cytosolic DAPs** | **Gene annotation** | **Primers** | **Sequences 5'-3'** |
| --- | --- | --- | --- | --- |
| 1. | *pecC* | *alr0525* | *pecC* F | GATGTTTGTGATTGAAGCGA |
|  |  |  | *pecC* R | CTATCTTTCCACCACGCTTA |
| 2. | *rps1* | *all0136* | *rps1 F* | TATAGAGAACAACTGCTGGC |
|  |  |  | *rps1 R* | GGATTTCCTCAACTGCTACA |
| 3. | *tufA* | *all4337* | *tufA* F | GATGGAGCAATTCTCGTAGT |
|  |  |  | *tufA* R | CGTCTTCCATCATGTCTTCT |
| 4. | *fus* | *all4338* | *fus* F | AATTATCCAAGGCTCTGCAA |
|  |  |  | *fus* R | CATCCGGTCTACGAGAATTT |
| 5. | *all4287* | *all4287* | *all4287*F | AAGACCCACAAAGTTCTCAA |
|  |  |  | *all4287*R | CTCTAGCAGACTCGAATGTT |
| 6. | *gap-2* | *all5062* | *gap-2* F | CGTCCGATTATCAAGGAACT |
|  |  |  | *gap-2* R | CTAATTCTGCCAAGTCGAGA |
| 7. | *fda* | *all4563* | *fda* F | GGTAATGCTCCTTCTACCTG |
|  |  |  | *fda* R | CTACATTTACAACTTCGCGG |
| 8. | *prk* | *alr4123* | *prk* F | TCACATTGTAGTGGTTGAGG |
|  |  |  | *prk* R | ACCTCTTTCTGCCATATCAC |
| 9. | *ctpA* | *all2500* | *ctpA* F | AGTAGAATTAACCGATGCCC |
|  |  |  | *ctpA* R | TTTGGCAGCAATCTGTTTAC |
| 10. | *cmpA* | *alr2877* | *cmpA* F | GAAGAATCACCCCGAAGAAT |
|  |  |  | *cmpA* R | AATTTCTGCCAGTTCTTTGC |
| 11. | *alr0803* | *alr0803* | *alr0803* F | ATCTGGTTTTGCGATGGATA |
|  |  |  | *alr0803* R | CCAGTAATAGCCCCTGTTTT |
| **Reference genes** | | | | |
| 12. | *rnpA* | *alr3413* | *rnpA* F | TACGCTCATTGGTGTCTCG |
|  |  |  | *rnpA* R | AACAACTGCTCTAATTCTTGC |
| 13. | *secA* | *alr4851* | *secA* F | AACCTACTACTACGACATCC |
|  |  |  | *secA* R | ACTTAATCACCTGTTCTTTCAA |

**Table S2** List of primers used for promoter analysis (EMSA)

| **S. N.** | **Genes**  **encoding cytosolic DAPs** | **Gene annotation** | **Primers** | **Sequences 5'**-**3'** |
| --- | --- | --- | --- | --- |
| 1. | *pecC* | *alr0525* | Pro *pecC* F | CGGAGCTCGTTAAGACAAAGCTTGT |
|  |  |  | Pro *pecC* R | GCGGTACCCATTTTGATAAACGCTC |
| 2. | *all4287* | *all4287* | Pro *all4287* F | CGGAGCTCCATTCCTCTAGTTAA |
|  |  |  | Pro *all4287* R | GCGGTACCGAGCCGATTATTAAA |
| 3. | *gap-2* | *all5062* | Pro *gap-2* F | CGGAGCTCTATTGCATATACCCT |
|  |  |  | Pro g*ap-2* R | GCGGTACCTATGCCTATAAGTTG |
| 4. | *prk* | *alr4123* | Pro *prk* F | CGGAGCTCTAGTGATTAGTGGAA |
|  |  |  | Pro *prk* R | GCGGTACCACAACCGAAATTAAC |
| 5. | *alr0803* | *alr0803* | Pro *alr0803* F | CGGAGCTCTTAAACTCGGAGATGTG |
|  |  |  | Pro *alr0803* R | GCGGTACCTGTAAAACTCGCCATGT |

**Table S3** List of spots showing change in abundance in recombinant *Anabaena* strains in response to γ-radiation (IR) or LexA overexpression

| **Samples** | **Change in abundance** | **Protein spot nos.** |
| --- | --- | --- |
| **AnpAM, IR^1*^** | Increased | 7, 8, 15, 17, 18, 19, 29, 30, 31, 39 |
| **An*lexA*^+^, IR^2*^** | Increased | 1, 7, 8, 9, 17, 28, 29, 34, 36, 39 |
| **AnpAM, IR^1*^** | Decreased | 4, 5, 6, 10, 11, 12, 13, 14, 16, 22, 24, 25, 26, 27, 28, 32, 37, 41 |
| **An*lexA*^+^, IR^2*^** | Decreased | 11, 12, 13, 14, 15, 18, 22, 23, 24, 25, 27, 32, 33, 37, 38 |
| **An*lexA*^+^, SIC^3#^** | Increased | 2, 3, 7, 8, 9, 10, 15, 17, 18, 19, 20, 21, 29, 30, 31, 33, 35, 36 |
| **An*lexA*^+^, SIC^3#^** | Decreased | 4, 5, 6, 12, 13, 14, 16, 22, 23, 24, 25, 28, 32, 38, 39, 40, 41 |

^#^SIC corresponds to sham-irradiated control cultures and **^*^**IR refers to γ-irradiated cultures.

**^1^** Proteome profile of AnpAM, IR samples are compared with that of AnpAM, SIC

**^2^** Proteome profile of An*lexA*^+^, IR samples are compared with that of An*lexA*^+^, SIC

**^3^** Proteome profile of An*lexA^+^*, SIC samples are compared with that of AnpAM, SIC

**Table S4** Quantification of band signal intensities in BN-PAGE gel using ImageJ software (http://rsb.info.nih.gov/ij) based on densitometric analysis.

| **Bands** | **Fold change in response to** | | |
| --- | --- | --- | --- |
|  | **LexA overexpression** | **γ-radiation (IR) in** | |
|  |  | **AnpAM** | **An*lexA*^+^** |
| **Band 1** | -1.59* | +1.18 | -1.38* |
| **Band 2** | -1.38* | +1.25 | -1.41* |
| **Band 3** | +1.24 | -1.05 | +1.05 |
| **Band 4** | +1.06 | +1.01 | -1.07 |
| **Band 5** | -1.17 | +1.23 | +1.11 |
| **Band 6** | +1.75* | +1.04 | -1.06 |
| **Band 7** | +1.31* | -1.06 | -1.02 |
| **Band 8** | +1.49* | -1.16 | -1.05 |
| **Band 9** | -1.13 | +1.57* | -1.31* |
| **Band 10** | -1.33* | +1.22 | +1.23 |

**Footnote**

The reported values are the mean fold change (based on three biological measurements) in the intensities of each band from AnpAM, IR and An*lexA*^+^, IR compared to AnpAM, SIC and An*lexA*^+^, SIC, respectively in order to determine the effect of γ-radiation, and that of An*lexA*^+^, SIC with AnpAM, SIC to determine the effect of LexA overexpression. SIC corresponds to sham-irradiated control cultures and IR refers to γ-irradiated cultures. The cut-offs of a fold change of 1.3 and *p* < 0.05 according to Student’s t-test with Tukey’s post-hoc comparison test.was applied to filter band intensities with significant differences and high confidence (differentially accumulated bands are indicated with asterisk

**Table S5** A comparison of the fold change in the intensities of 41 DAP spots detected in this study due to LexA overexpression (AnpAM, SIC *vs* An*lexA*^+^, SIC), in which both recombinant *Anabaena* strains, AnpAM (vector control) and An*lexA*^+^ (LexA-overexpressing) were kept in dark, with the same spots detected in an earlier 2D-PAGE analysis in response to LexA overexpression (Kumar et al., 2018), in which both strains were grown in continuous light. Significant up- and down-accumulation in DAP spots (*p* < 0.05) are shown by green and red colors, respectively.

| **Spot No.** | **Protein name** | **Fold change in response to LexA overexpression** | |
| --- | --- | --- | --- |
|  |  | **Current study** | **Kumar et al. (2018)*** |
| 23 | C-phycocyanin subunit beta | **-4.26** | **-8.17** |
| 24 |  | **-4.66** | **-12.13** |
| 1 | Phycobilisome 32.1 kDa linker polypeptide, phycocyanin-associated, rod | **1.26** | **-1.28** |
| 2 |  | **2.69** | **Not detected in both strains** |
| 3 |  | **3.34** | **3.16** |
| 5 |  | **< -10** | **Not detected in both strains** |
| 11 |  | **-1.02** | **Not detected in both strains** |
| 10 | Phycobilisome rod-core linker polypeptide | **1.71** | **2.11** |
| 4 | Phycobiliprotein ApcE | **< -10** | **-2.53** |
| 12 | Phycobilisome 34.5 kDa linker polypeptide, phycoerythrocyanin-associated | **-3.12** | **-2.67** |
| 13 |  | **-2.58** | **-2.28** |
| 14 |  | **-5.76** | **-1.68** |
| 17 | Flavodoxin | **3.13** | **Not detected in both strains** |
| 7 | Thioredoxin 2 | **2.43** | **-1.33** |
| 25 | Peroxiredoxin | **-1.51** | **-1.74** |
| 37 | 30S ribosomal protein S1 | **1.36** | **1.15** |
| 6 | 30S ribosomal protein S8 | **-1.85** | **Not detected in both strains** |
| 22 | Elongation factor Tu | **-5.00** | **-1.29** |
| 38 | Elongation factor G | **-3.03** | **-2.11** |
| 18 | Peptidyl-prolyl cis-trans isomerase B | **2.56** | **Not detected in both strains** |
| 20 | Enoyl-[acyl-carrier-protein] reductase [NADH] | **> 10** | **> 10** |
| 21 | Ribulose-phosphate 3-epimerase | **> 10** | **> 10** |
| 26 | Glyceraldehyde-3-phosphate dehydrogenase 2 | **1.03** | **1.13** |
| 27 |  | **-1.32** | **1.22** |
| 28 |  | **-2.27** | **1.57** |
| 29 | Fructose-bisphosphatealdolase | **2.21** | **1.56** |
| 30 |  | **1.69** | **-1.40** |
| 31 |  | **2.22** | **< -10** |
| 36 |  | **3.86** | **1.13** |
| 32 | Phosphoribulokinase | **-2.70** | **Not detected in both strains** |
| 35 | Acetyl CoenzymeA-synthetase | **1.91** | **1.18** |
| 33 | Carboxyl-terminal protease | **> 10** | **Not detected in both strains** |
| 8 | Putative pterin-4a-carbinolamine dehydratase | **1.77** | **2.87** |
| 34 | Bicarbonate transport bicarbonate-binding protein | **-1.19** | **1.02** |
| 39 | Cystathionine beta-synthase | **-2.04** | **1.19** |
| 40 | Nutrient stress-induced DNA-binding protein | **< -10** | **-1.68** |
| 41 | DNA-directed RNA polymerase subunit beta' | **< -10** | **1.57** |
| 9 | All2133 (Hypothetical) | **4.19** | **-1.30** |
| 15 | Alr0803 (Hypothetical) | **3.72** | **1.29** |
| 16 | Alr1144 (Unknown protein) | **-3.38** | **-7.31** |
| 19 | Alr0882 (Hypothetical) | **1.94** | **Not detected in both strains** |

**Footnote**

* Kumar, A., Kirti, A., Rajaram, H., 2018. Regulation of multiple abiotic stress tolerance by LexA in the cyanobacterium *Anabaena* sp. strain PCC7120. Biochim. Biophys. Acta-Gene Regul. Mech. 1861, 864-877.
